# Pulmonary infection interrupts acute cutaneous wound healing through disruption of chemokine signals

**DOI:** 10.1101/2020.05.08.084442

**Authors:** Meredith J. Crane, Yun Xu, Sean F. Monaghan, Benjamin M. Hall, Jorge E. Albina, William L. Henry, Holly L. Tran, Karisma R. P. Chhabria, Alexander R. D. Jordon, Lindsey Carlsen, Amanda M. Jamieson

**Affiliations:** Division of Biology and Medicine, Department of Molecular Microbiology and Immunology, Brown University, Providence, Rhode Island, United States; Division of Surgical Research, Department of Surgery, Rhode Island Hospital and the Warren Alpert School of Medicine of Brown University, Providence, Rhode Island, United States

**Keywords:** Wound healing, innate immune response, chemokine, immune cell trafficking, lung infection

## Abstract

Studies of the immune response typically focus on single-insult systems, with little known about how multi-insult encounters are managed. Pneumonia in patients recovering from surgery is a clinical situation that exemplifies the need for the patient to mount two distinct immune responses. Examining this, we have determined that poor wound healing is an unreported complication of pneumonia in laparotomy patients. Using mouse models, we found that lung infection suppressed the trafficking of innate leukocytes to wounded skin, while pulmonary resistance to the bacterial infection was maintained. The dual insults caused distinct systemic and local changes to the inflammatory response, the most striking being a rapid and sustained decrease in chemokine levels at the wound site of mice with pneumonia. Remarkably, replenishing wound chemokine levels completely rescued the wound-healing rate in mice with a pulmonary infection. These findings have broad implications for understanding the mechanisms guiding the innate immune system to prioritize inflammatory sites.

**One Sentence Summary:** Chemokine-mediated signaling drives the prioritization of innate immune responses to bacterial pulmonary infection over cutaneous wound healing.

**Highlights:**

- Human laparotomy patients with pneumonia have an increased rate of incision dehiscence, and this observation can be recapitulated in mouse models of bacterial lung infections and skin wounds.
- Lung infection causes rapid and sustained suppression of skin wound chemokine and inflammatory cytokine production as well as leukocyte recruitment.
- Unique systemic shifts in the immune compartment occur with two inflammatory insults, including the cytokine/chemokine signature and the mobilization, recruitment, and phenotype of innate leukocytes.
- Restoration of chemokine signaling in the wounds of mice that have a lung infection results in increased neutrophil trafficking to the wound site and rescues the rate of healing.

**Graphical Abstract:** 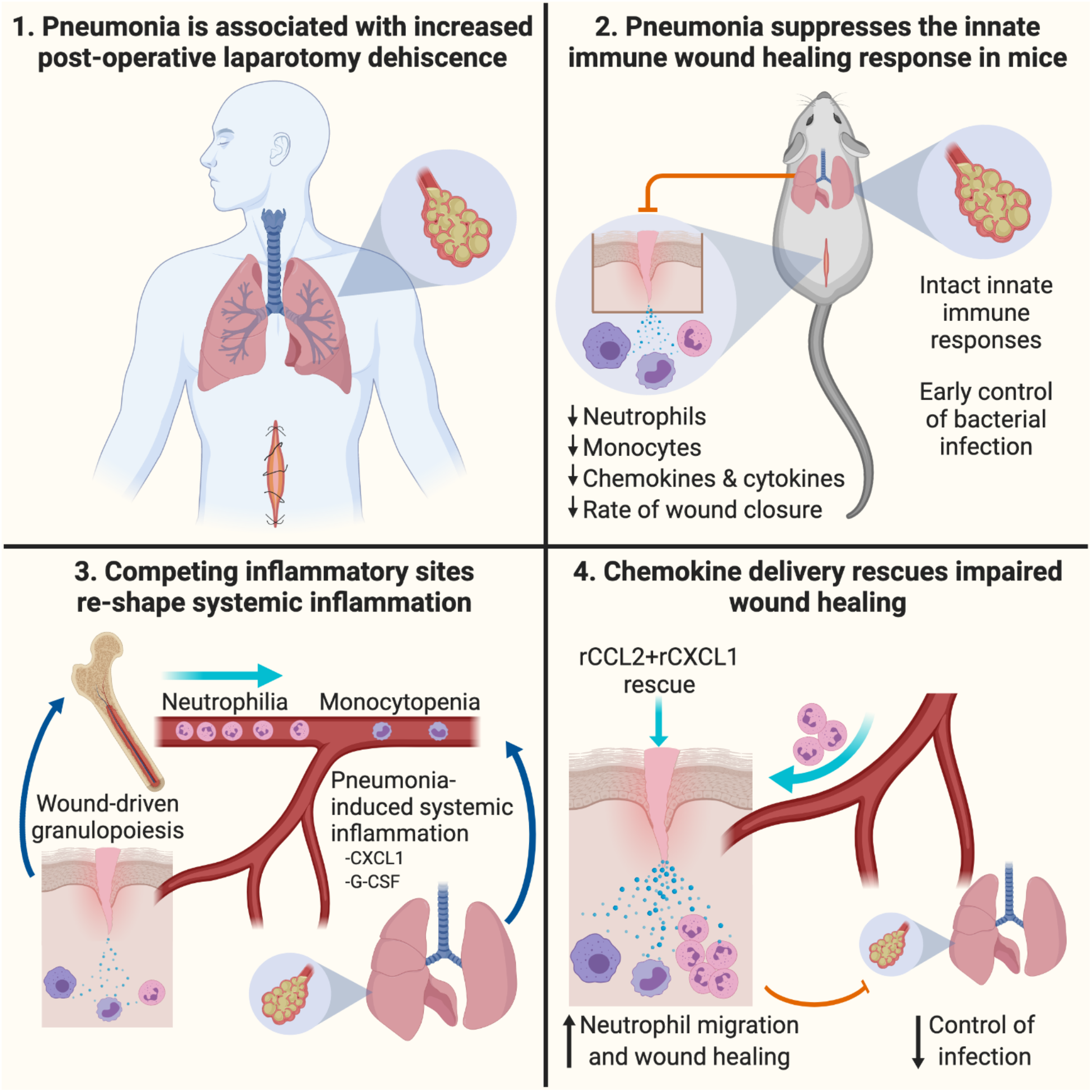

## Introduction

The immune system is essential in many processes that are required to maintain health, ranging from host-defense, disease tolerance, and the response to cancer, to development, neurophysiology, metabolism, and tissue repair (*1*–*5*). Many of these responses must occur simultaneously. However, we have a limited understanding of how multi-insult encounters are managed by the innate immune system, as studies of immune function have classically focused on single-insult encounters, such as infection or injury. We propose that when faced with more than one insult, a form of immune triage occurs, where the most life-threatening risk receives the attention needed to resolve the threat. To examine this, we turned to the example of hospitalized surgical patients.

Post-operative patients in the hospital are at risk of developing secondary healthcare-associated infections, such as pneumonia, and this occurrence extends the length of hospital stay and increases the rate of morbidity and mortality (*6*–*8*). Patients with traumatic or surgical injuries rely on intact innate immune responses to drive acute wound healing (*9*–*23*). However, the innate immune response is also essential in preventing and clearing infections, raising the possibility that there will be increased stress on the immune response during the post-operative recovery period if complicated by a concurrent infection (*9*–*23*). In cases of post-operative pneumonia, it is not known how these inflammatory sites interact and shape the immune response, and how this may affect the ability to successfully heal a wound. With a specific focus on the components of the innate immune system that orchestrate the acute phases of wound healing, including neutrophils, inflammatory monocytes, macrophages, and an array of inflammatory cytokines and chemokines (*24*), we hypothesized that the cellular innate immune response is insufficient to meet the demands of these dual insults, resulting in the prioritization of one inflammatory site.

In order to address our hypothesis, we assessed surgical patient data and established murine models of post-operative pulmonary infection. We report here a previously unknown increased incidence of poor wound healing in surgical patients who develop pneumonia. To understand this at a mechanistic level, we developed mouse models of pulmonary infection following surgical wounding, focusing on bacterial lung infection because nosocomial pneumonia is most commonly caused by a variety of bacterial species (*25*). Recapitulating patient data, lung infection caused delayed cutaneous wound healing in mice. Inflammatory cellular and cytokine responses in the wound were rapidly suppressed by lung infection, while effects of the two inflamed sites generated a unique immune signature in the bone marrow and blood. A loss of chemokine signals was the underlying cause of decreased wound cellularity and poor healing in lung-infected mice, as the therapeutic application of monocyte and neutrophil chemoattractants to the wound site fully restored healing. Our findings introduce a mechanistic basis for how the immune system uses chemokine networks to triage, or prioritize, inflammatory sites.

## Materials and Methods

### Analysis of surgical patient data

The American College of Surgeons (ACS) National Surgical Quality Improvement Program (NSQIP) Participant Use Data File (PUF) for 2015 was utilized in this study, 2015 being the most recent available dataset. The ACS NSQIP PUF is a HIPAA compliant file with no protected health information that was accessed after approval from ACS NSQIP. For 2015 there were over 885,000 operative cases from 603 hospitals. Trained nurses enter all data in the NSQIP database with the focus on quality improvement, therefore complications like dehiscence are less likely to be missed as would be expected in self-reporting situations. All patients with a primary CPT code involving a laparotomy were included for analysis, regardless of emergent status. These patients were assessed for a dehiscence after pneumonia based on the postoperative days reported. Age was compared based upon two groups: 18-40 and >65. This was done to have a buffer age range (41–64) that would show a true physiologic difference based upon age.

### Mice

All animal studies were approved by the Brown University Institutional Animal Care and Use Committee and carried out in accordance with the Guide for the Care and Use of Animals of the National Institutes of Health. The animal protocol (number 1608000222) was approved on September 26, 2016, and the annual continuation of this protocol was approved on September 28, 2018. C57BL/6J mice were purchased from The Jackson Laboratory. B6.SJL-*Ptprc^a^Pepc^b^*/BoyJ (CD45.1 congenic) mice were bred in-house. Male mice 8-12 weeks of age were used in all experiments.

### Polyvinyl alcohol sponge implantation

Prior to surgeries, mice received intraperitoneal injections of ketamine (60-80 mg/kg) and xylazine (30-40 mg/kg) to induce anesthesia and analgesia. The dorsum was shaved and cleaned with povidone-iodine solution and isopropyl alcohol. Under sterile conditions, six 1cm×1cm×0.3cm sterile PVA sponges (Ivalon, PVA Unlimited, Inc.) were placed into subcutaneous pockets through a 2cm midline dorsal incision. The incision was then closed with surgical clips.

### Full-thickness tail wounding

Prior to surgery the tail was cleaned with povidone-iodine solution and isopropyl alcohol. Using a scalpel, a 1cm × 0.3cm area of the skin was excised 0.5cm from the base of the tail. The wound bed was then covered with a spray barrier film (Cavilon, 3M). Length and width measurements were taken at the midpoints of the wound bed using calipers, and these measurements were used to calculate the wound area. A secondary measurement of wound area on photographed wounds was analyzed using ImageJ software (NIH). Tail wound images were acquired from a fixed position using a 12-megapixel iSight camera. All measurements were done in a blinded fashion.

### Bacterial pulmonary infection

Mice were anesthetized by intraperitoneal injection of ketamine (60-80 mg/kg) and xylazine (30-40 mg/kg). Mice were given 2×10^7^ CFU *Klebsiella oxytoca* or 5×10^6^ CFU *Streptococcus pneumoniae* intranasally in a volume of 30μL, with sterile saline as the vehicle.

### Wound fluid and cell isolation

Mice were euthanized by CO_2_ asphyxiation. To collect wound fluid, three sponges isolated from the right of midline from each animal were placed in the barrel of a 5mL syringe and centrifuged in a collection tube. For cell collection, three sponges from the left of midline from each animal were placed in 1x HBSS medium (1% FBS/penicillin/streptomycin/1M HEPES) and cells were isolated by mechanical disruption using a Stomacher (Tekmar). Wound cells were washed with 1x HBSS medium and red blood cells lysed. Cell counts were obtained using a Moxi Z Automated Cell Counter (Orflo) or an Attune NxT flow cytometer (ThermoFisher).

### Plasma and blood cell collection

Blood was collected retro-orbitally in the presence of heparin. Plasma was separated from red blood cells and leukocytes by centrifugation in Wintrobe Tubes (CMSLabcraft). Leukocytes were contained within the buffy coat layer at the interface of plasma and red blood cells. Residual red blood cells in the buffy coat layer were removed by lysis. Cells were counted using a Moxi Z Automated Cell Counter (Orflo) or an Attune NxT Flow Cytometer (ThermoFisher).

### Bronchoalveolar lavage and lung cell preparation

To collect bronchoalveolar lavage fluid (BALF), a BD Venflon IV catheter was inserted into the exposed trachea. The catheter was used to flush the bronchoalveolar space twice with 1ml of sterile 1xPBS. Cell-free supernatants were collected for cytokine analyses and protein content quantification. Cells were counted with a Moxi Z Automated Cell Counter (Orflo) or an Attune NxT Flow Cytometer (ThermoFisher).

To isolate cells from lung tissue, the right superior and middle lobes were perfused with 20 ml of PBS then minced. The tissue was incubated for 45 minutes at 37° C in DMEM containing type 4 collagenase (Worthington Biochemical Corporation) and DNAse I (Sigma-Aldrich). The digested tissue was strained at 70μM. The cell pellet was then re-suspended in 4ml of 40% Percoll/PBS and carefully layered over 4ml of 80% Percoll/PBS. The gradient was centrifuged at room temperature for 20 minutes at 600g with low acceleration and deceleration. Cells at the Percoll gradient interface were collected and washed once with 10ml PBS containing 5% FBS.

### Pulmonary CFU analysis

The right superior lung lobe was homogenized in sterile 1x PBS. Serial dilutions of homogenates were plated onto Trypticase Soy Agar with 5% Sheep Blood (TSA II, BD) for quantitation of colony forming units (CFU).

### Quantitation of BALF total protein

The bicinchoninic (BCA) assay was used to measure the concentration of protein in the BALF according to manufacturer instructions (Pierce Chemical Co.). Each sample was tested against an albumin standard.

### Flow cytometry analysis of cell subsets

#### The following antibodies were used to identify cell subsets

Ly6C-FITC (AL-21, BD Biosciences), F4/80-APC eFluor660 (BM8, eBioscience), Siglec-F-PE or AR700 (E50-2440, BD Biosciences), CD11c-PE or BV711 (HL3, BioLegend), Ly6G-PerCP eFluor710 or V450 (1A8, eBioscience or BD Biosciences), CD45.2-APC/Fire750 or V450 (104, BioLegend or eBioscience), CD45.1-PE (A20, eBioscience), and CD11c-BV711 (N418, BioLegend). Dead cells were excluded from analyses using Fixable Viability Dye APC BV506 (eBioscience).

#### Surface staining

Cells were treated with anti-CD16/CD32 Fc receptor blocking antibody (clone 2.4G2) in 1x PBS (1% FBS) for 10 minutes on ice. Cells were then centrifuged and resuspended in 1x PBS (1% FBS) containing antibodies and incubated for 15 minutes on ice. Cells were washed with 1x PBS then incubated with Fixable Viability Dye diluted in 1x PBS for 15 minutes on ice. Cells were washed, then fixed with 1% paraformaldehyde for 15 minutes on ice.

Samples were acquired using an Attune NxT Acoustic Focusing Cytometer with Attune Software. Analyses were performed using FlowJo v10 software (Tree Star, Inc.). Gate placement was determined using fluorescence minus one and unstained control samples.

### Cytokine analysis

The concentration of cytokines and chemokines, with the exception of G-CSF, CXCL1 and CXCL5, in wound fluid, plasma, and BALF was measured using a custom LEGENDplex bead-based immunoassay (BioLegend) according to manufacturer instructions. G-CSF, CXCL1 and CXCL5 concentrations were determined using DuoSet sandwich ELISA kits (R&D Systems) according to manufacturer instructions.

### Bone marrow cell isolation and adoptive transfer

For immunophenotyping, femurs were collected from C57BL/6J mice in 1x HBSS. For cell adoptive trasnfers, femurs and tibias were collected from CD45.1 congenic mice in sterile 1x HBSS. Bone marrow was collected from femurs and/or tibias by flushing with sterile 1x HBSS medium (1% FBS/penicillin/streptomycin/1M HEPES) using a syringe and red blood cells were lysed with water under sterile conditions. Isolated cells were counted with a Moxi Z Automated Cell Counter (Orflo) or an Attune NxT Flow Cytometer (ThermoFisher). For adoptive transfer, 5×10^6^ bone marrow cells cells in a volume of 100μL of sterile 1x PBS were transferred retro-orbitally to recipient wild-type C57BL/6J mice.

### Application of exogenous chemokines to tail wounds

Fibrin sealant (Tisseel, Baxter) was delivered to the wound bed by co-application of thrombin and fibrinogen, which were prepared under sterile conditions according to the manufacturer instructions. The sealer protein protease inhibitor was omitted from the mixture to facilitate delivery of the recombinant chemokines to the wound bed via fibrin degradation. Directly before application, recombinant murine CCL2 (Peprotech) and recombinant murine CXCL1 (Peprotech) were mixed into the fibrinogen component. Control mice were treated with Tisseel without chemokines. The fibrinogen and thrombin components were maintained at 37°C to avoid polymerization. Using two pipets, equal volumes of fibrinogen and thrombin were simultaneously applied to the wound beds of anesthetized mice and allowed to polymerize. Treatments were given every day from wound days 1 to 7, then every other day for the remainder of the experiment. Each chemokine treatment contained 10ng of recombinant CCL2 and 10ng of recombinant CXCL1. Application volumes were adjusted according to wound bed area and ranged from 30uL to 10uL.

### Application of exogenous chemokines to PVA sponge wounds

Recombinant murine CCL2 (Peprotech) and recombinant murine CXCL1 (Peprotech) were diluted in 1x PBS for injection into implanted PVA sponges. The backs of mice were cleaned with iodine solution and isopropyl alcohol. 0.5ug of each chemokine mixed in a total volume of 50uL of PBS was injected through the skin and into the center of each sponge for a total treatment of 3ug per wound. Control mice received injections of PBS vehicle. Mice were injected on wound days 5 and 6, and sponges were isolated on wound day 7.

### Statistical analysis

Biostatistical analyses of murine samples were carried out using the GraphPad Prism software package. For comparison of two groups the nonparametric Mann Whitney test was used. To compare 3 or more groups the Kruskal-Wallis one-way analysis of variance or, for data sets with multiple time points, a two-way analysis of variance with Tukey’s multiple comparisons test was used. All the groups were compared to each other. For clarity in murine experiments, only statistically significant differences between wound + *K. oxytoca* and control, wound, or *K. oxytoca* groups are presented. Unless otherwise noted, statistically significant changes between control and wound + *K. oxytoca* are denoted by *, between wound and wound + *K. oxytoca* are denoted by %, and between *K. oxytoca* and wound + *K. oxytoca* are denoted by #. Differences were considered significant if the p value was calculated to be ≤ 0.05. Clinical data were managed and analyzed using SAS (Cary, NC) using the included generalized linear mixed model and alpha was set to 0.05.

## Results

### Pneumonia is associated with wound dehiscence among patients with abdominal incisions

While it is well known that pneumonia is a risk of hospitalization, especially in post-surgical and trauma patients, it is not known how pneumonia impacts the ability to heal a wound (*26*–*30*). To address this question, we consulted the American College of Surgeons (ACS) National Surgical Quality Improvement Program (NSQIP) Participant Use Data File (PUF) to assess the rate of abdominal incision dehiscence among patients with or without pneumonia. Dehiscence is a post-surgical complication in which the wound ruptures along the site of the incision, and it is a clear indicator of a poorly healing surgical wound. Of over 885,000 cases in the ACS NSQIP PUF for 2015, 89,608 cases were included as they had a midline abdominal incision. A total of 1221 patients had a dehiscence (1.4%). When assessing patients who had a dehiscence and pneumonia, the dehiscence rate was 3.16%, compared to a dehiscence rate of 1.28% among patients who did not have pneumonia (p<0.0001, Table 1). Surgical site infection is typically associated with an increased risk of dehiscence; however, pneumonia did not make it more or less likely to have a surgical site infection (6.3% vs 6.7%, p=0.2829, Supplementary Table 1). Age has also been associated with increased dehiscence, and in a model to predict dehiscence, both age >65 (F=8.4, p=0.00037) and pneumonia (F=59.57, p<0.0001) were significant, but the interaction between the two was not (F=0.51, p=0.4749). This increased wound dehiscence was also specific for lung infections as another common hospital-acquired infection, urinary tract infection, did not increase the rate of dehiscence (Supplemental Table 2). This analysis demonstrates that the onset of pneumonia in patients with surgical injuries is a risk factor for complications in wound healing.

**Table 1.** Rate of abdominal wound dehiscence among surgical patients with pneumonia

| PNEUMONIA | DEHISCENCE |  | TOTAL |
| --- | --- | --- | --- |
|  | No Dehiscence (% of Total) | Dehiscence (% of Total) |  |
| No Pneumonia | 84774 (98.72%)<sup>^</sup> | 1103 (1.28%)<sup>^</sup> | 85877 |
| Pneumonia | 3613 (96.84%)<sup>#</sup> | 118 (3.16%)<sup>#*</sup> | 3731 |

<sup>^</sup>Percent of total number of patients in "No Pneumonia" group
<sup>#</sup>Percent of total number of patients in "Pneumonia" group
\* $p \leq 0.0001$ versus % dehiscence in "No Pneumonia" group

### Excisional tail skin wound closure is delayed in mice with lung infection

We established murine models to determine the mechanisms of impaired wound healing in pneumonia patients. To assess the rate of wound closure, a 1cm × 0.3cm excisional tail skin wound was employed. Tail skin was chosen as the site of wounding because it is firm, lacks fur, and relies primarily on re-epithelialization, which is more akin to human skin healing than other murine models of wound closure (*31*–*33*). Initial wound area measurements were obtained on day 1 post-wounding. At this time, mice either remained uninfected or were infected intranasally with the Gram-negative opportunistic bacterium *Klebsiella oxytoca* (wound + *K. oxytoca*) (*34*, *35*). The wound area was measured over the course of 15 days, and it is reported as a fraction of the day 1 wound area (Figure 1a). From days 7 to 15 post-wounding, wound + *K. oxytoca* mice had significantly larger wounds compared to wounded mice alone (Figure 1b, Figure S1). These data indicate that pulmonary infection causes a delay in tail wound closure.

**Fig. 1.**
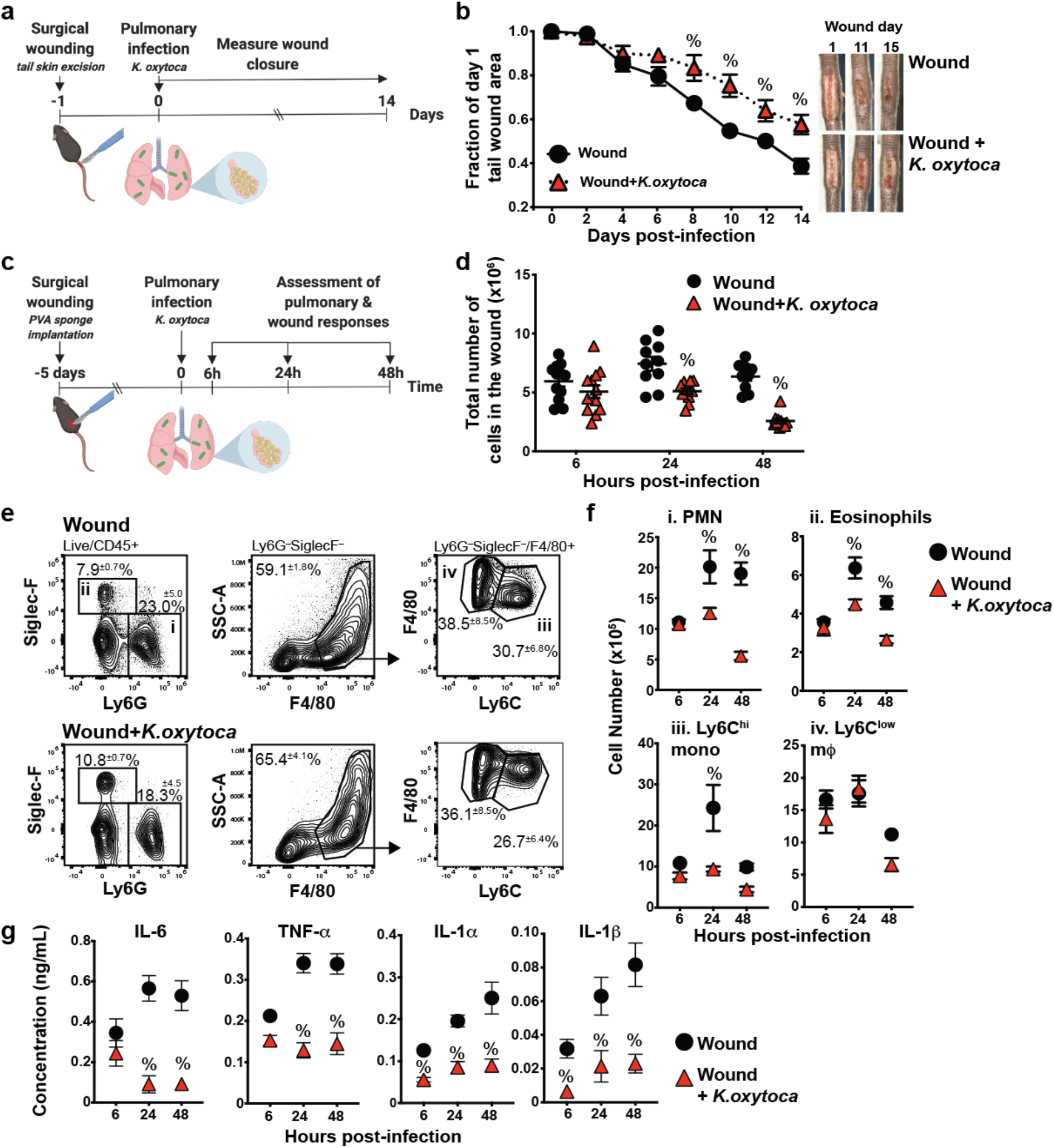
Pulmonary *K. oxytoca* infection impairs wound healing. **a)** To determine the effect of lung infection on wound closure, excisional tail wounds were performed on C57BL/6J mice (wound), and a cohort was infected intranasally with *K. oxytoca* (wound + *K. oxytoca*) on wound day 1. The tail wound area was measured every other day beginning on day 1. **b)** Tail wound closure was measured in mice infected with *K. oxytoca* infection and compared to uninfected mice. **c)** The PVA sponge wound model was used to assess the effects of pulmonary infection on cellular wound healing responses. Sponges were surgically implanted, and a cohort was infected intranasally with *K. oxytoca* (wound + *K. oxytoca*) 5 days later. **d)** Wound cellularity was assessed 6, 24, and 48h post-infection. **e)** Flow cytometry analysis of wound cells isolated from uninfected or infected mice 48h post-infection shows the frequency of Ly6G^+^ neutrophils (PMN, i), Siglec-F^+^ eosinophils (ii), F4/80^+^Ly6C^hi^ monocytes (mono, iii), and F4/80^+^Ly6C^low^ macrophages (mΦ, iv). **f)** The absolute number of innate leukocyte populations in wounds of uninfected of *K. oxytoca*-infected mice. **g)** Wound fluids were assayed by LegendPlex for the proinflammatory cytokines IL-6, TNF-α, IL-1α, and IL-1β. Data are shown as the mean±SEM with minimum n=10 mice per group from three independent experiments. Results are considered statistically significant when p ≤ 0.05. Statistically significant changes between wound and wound + *K. oxytoca* are denoted by %.

### Pulmonary *K. oxytoca* infection decreases innate leukocyte cellularity in the wound

Activation of the innate immune system is essential for the early stages of wound healing. To investigate the effects of pulmonary infection on the acute wound healing response, mice were wounded by the dorsal subcutaneous implantation of polyvinyl alcohol (PVA) sponges. This model follows the acute stages of wound healing, and allows for the retrieval of cells and fluids from the implanted sponges after their removal (*10*, *11*, *16*, *20*, *33*, *36*). By 7 days post-sponge implantation, a large number of leukocytes are recoverable from implanted sponges, and the cytokine milieu reflects the transition from the inflammatory to the repair phases of wound healing (*10*, *11*, *16*, *20*, *36*).

Mice with PVA sponge wounds remained uninfected or were infected intranasally with *K. oxytoca* (wound + *K. oxytoca*) five days after sponge implantation in order to synchronize the inflammatory response to infection (Figure S2) with an established acute wound healing leukocyte response (*10*). Wound cellularity was assessed 6, 24, and 48 hours after infection (Figure 1c). Beginning at 24 hours post-infection, fewer infiltrating immune cells were isolated from sponges removed from wound + *K. oxytoca* mice compared to wounded mice alone (Figure 1d). Initiating *K. oxytoca* infection in wounded mice on wound day 1 similarly led to a decrease in wound cellularity at 24- and 48-hours post-infection (Figure S3), indicating that this suppression was not specific to the timing of infection after wounding.

The wound leukocyte milieu in uninfected and *K. oxytoca*-infected mice was assessed to determine whether pulmonary infection altered a specific cell type in the wound. Mice were wounded by PVA sponge implantation, and a cohort was infected intranasally with *K. oxytoca* on wound day 5 (Figure 1c). Cell populations in the wound were identified by flow cytometry analysis 6, 24, and 48 hours post-infection. CD45^+^ innate leukocytes are the predominant cell type in PVA sponge wounds at these times, and consist primarily of Ly6G^+^ neutrophils, Siglec-F^+^ eosinophils, F4/80^+^Ly6C^hi^ monocytes, and F4/80^+^Ly6C^low^ monocyte-derived macrophages (*10*, *11*, *20*). Representative gating of wound cells 48 hours after infection (wound day 7) is shown in Figure 1e, and the full gating strategy to identify these cell populations is reported in Figure S4. The relative percentage of neutrophils, monocytes, macrophages, and eosinophils was the same in wound and wound + *K. oxytoca* groups (Figure 1e). In contrast, the absolute number of all innate leukocyte populations examined was lower in wound + *K. oxytoca* mice than wounded mice alone at 24- and 48-hours post-infection, with the exception of F4/80^+^Ly6C^low^ macrophages (Figure 1f). This indicates that the cellularity defect lies primarily with cells that migrate to the wound from the circulation (*37*). Together, these data demonstrate that pulmonary infection suppresses the number of innate leukocytes in the wound, which results in an overall loss of wound cellularity.

### Wound cytokines are suppressed in *K. oxytoca*-infected mice

Coordinated wound cytokine responses are necessary for the normal progression of the repair response (*10*–*12*, *16*, *17*, *36*, *38*). The effect of pulmonary infection on a time course of wound inflammatory cytokine concentrations was assessed in PVA sponge wound fluids from mice that were treated as shown in Figure 1c. As expected, the cytokines TNF-α, IL-6, IL-1β and IL-1α were present in the wound fluid at all time points examined. IL-1β and IL-1α were suppressed in wound fluids as soon as 6 hours post-infection. By 24 hours post-infection, wound fluids had lower concentrations of all cytokines compared to uninfected mice (Figure 1g), demonstrating that pulmonary infection causes suppression of cytokine responses in PVA sponge wounds.

### Prior wounding alters the kinetics of pulmonary cytokine and cellular responses but does not impact resistance to *K. oxytoca* infection

In response to *K. oxytoca* infection, inflammatory cytokines and chemokines are rapidly produced in the lung (*21*). To determine whether an ongoing wound healing response influenced the ability to respond to a pulmonary bacterial infection, IL-6, TNF-α, IL-1α, IL-1β, and GM-CSF levels in the bronchoalveolar lavage fluid (BALF) were assessed in control, wound, *K. oxytoca*, or wound + *K. oxytoca* groups (Figure 2a). Mice were wounded and/or infected as previously described (Figure 1c). *K. oxytoca* infection alone induced all cytokines as early as 6 hours after infection. At 6 hours post-infection, wound + *K. oxytoca* mice had a significantly higher concentration of IL-6 in the BALF than infected mice alone. In contrast, the production of IL-1β and GM-CSF in wound + *K. oxytoca* mice was delayed compared to infected mice alone. These data indicate that the presence of a wound alters the initial cytokine response to pulmonary infection.

**Fig. 2.**
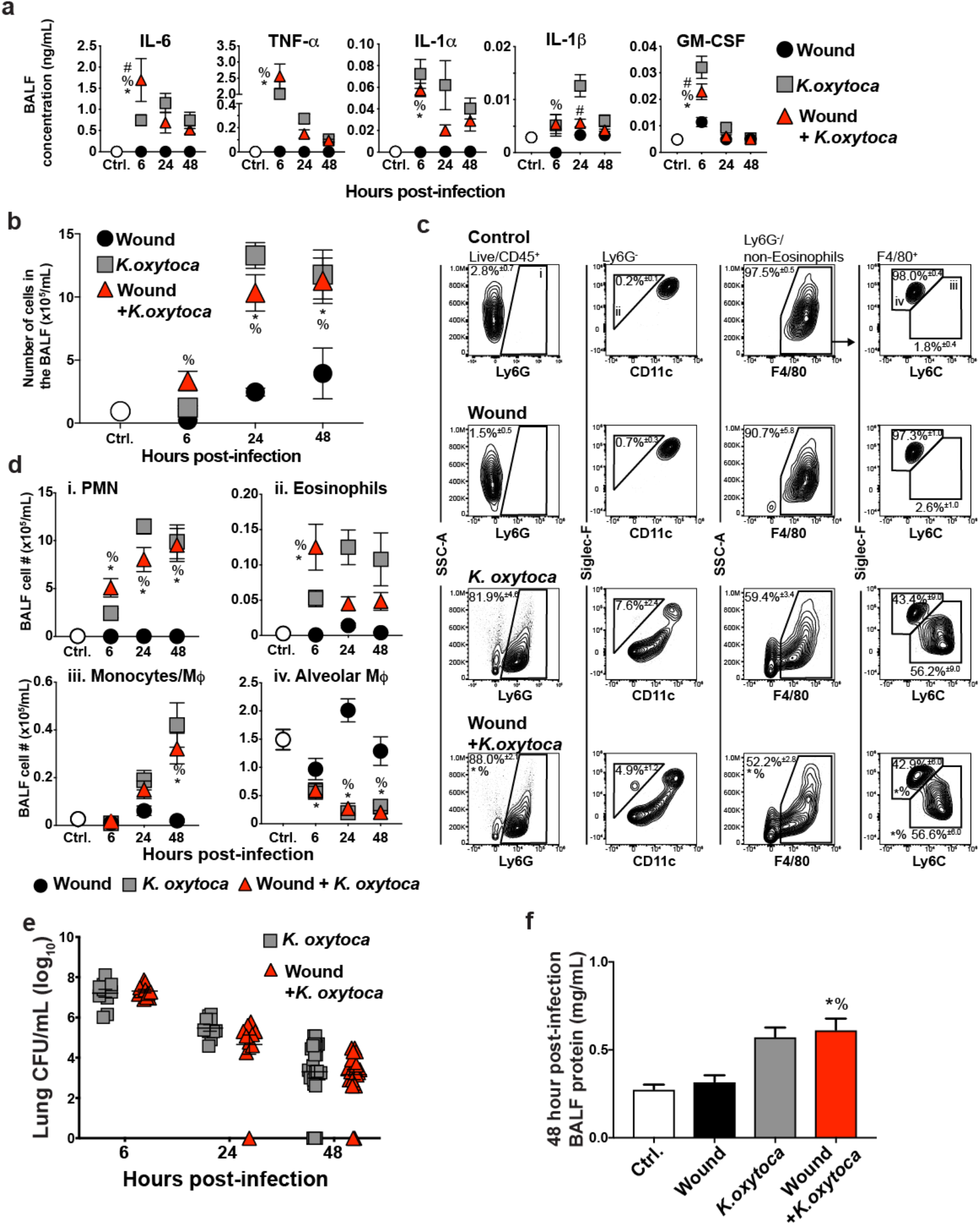
PVA sponge wounds alter the pulmonary cellular and cytokine response but not early control of *K. oxytoca* infection. C57BL/6J mice were wounded by PVA sponge infection (wound), and a cohort was infected 5 days later (wound + *K. oxytoca*). Unwounded mice were also infected with *K. oxytoca* (*K. oxytoca*) or remained uninfected (ctrl.). **a)** The BALF was assayed with LegendPlex for a panel of cytokines that are induced in response to bacterial infection. **b)** A time course of BALF cellular content was determined for all experimental groups. **c)** Flow cytometry analysis shows the proportion of BALF Ly6G^+^ neutrophils (PMN, i), CD11c^−^Siglec-F^+^ eosinophils (ii), F4/80^+^Ly6C^hi^Siglec-F^−^ monocytes/macrophages (mΦ) (iii), and F4/80^+^Ly6C^low^Siglec-F^+^ alveolar macrophages (mΦ) (iv) at 48h post-infection. **d)** The absolute number of BALF leukocyte populations over time. **e)** Lung *K. oxytoca* titers were determined in *K. oxytoca*-infected and wound + *K. oxytoca* mice. **f)** The BALF protein content at 48h post-infection was measured by BCA assay to assess pulmonary vascular permeability. Data are shown as the mean±SEM with a minimum n=10 mice per group from three independent experiments. Results are considered statistically significant when p ≤ 0.05. Statistically significant changes between control and wound + *K. oxytoca* are denoted by *, between wound and wound + *K. oxytoca* are denoted by %, and between *K. oxytoca* and wound + *K. oxytoca* are denoted by #.

To determine if the observed changes in BALF cytokine production in wound + *K. oxytoca* mice influenced the ability to mount a cellular response to the bacterial pathogen, we examined leukocyte populations in the BALF. *K. oxytoca* infection caused an increase in BALF cellularity compared to control mice. A similar increase in BALF cellularity was observed in wound + *K. oxytoca* mice (Figure 2b). The distribution of CD45^+^ innate leukocytes in the BALF of control, wound, *K. oxytoca*, and wound + *K. oxytoca* groups was also determined at 6, 24, and 48 hours post-infection. Representative flow cytometry analyses from 48 hours post-infection are shown in Figure 2c, and the full BALF gating strategy is presented in Figure S5. Nearly all cells isolated from the BALF of uninfected mice were F4/80^+^Ly6C^low^Siglec-F^+^ alveolar macrophages, while Ly6G^+^ neutrophils were the predominant innate leukocyte population in *K. oxytoca*-infected mice. F4/80^+^Siglec-F^low^Ly6C^hi^ monocytes/macrophages also accumulated in the BALF of infected mice. Wounding did not significantly alter the kinetics of the lung-infiltrating leukocytes examined in infected mice, although the number of Ly6G^+^ neutrophils was modestly elevated in the BALF of wound + *K. oxytoca* mice at 6 hours post-infection (Figure 2d).

To determine if the presence of a PVA sponge wound impacted the ability to control the pulmonary infection, bacterial titers were measured in *K. oxytoca* and wound + *K. oxytoca* groups. *K. oxytoca* titers were the same in infected mice with or without wounds at all time points examined (Figure 2e). BALF protein content was also measured at 48 hours post-infection to assess pulmonary vascular permeability in all experimental groups. *K. oxytoca* and wound + *K. oxytoca* mice had similar increases in BALF protein content after infection compared to uninfected groups (Figure 2f). Overall, these data show that the presence of a wound primes the pulmonary environment for rapid induction of IL-6 and neutrophil responses; however, this does not impact the ability to respond to the lung infection or cause excess vascular permeability.

To determine whether the suppressive effect of *K. oxytoca* infection on dermal wound healing was pathogen-specific, or more broadly applicable to other bacterial pulmonary infections, the experiments described in Figure 1c were repeated using pulmonary infection with the Gram-positive bacterium *Streptococcus pneumoniae*. As infection with *S. pneumoniae* is lethal after several days, we focused on the innate immune wound healing response using the PVA sponge wound model rather than the excisional tail skin wound model (Figure S6a). Similar to what was observed with *K. oxytoca* infection, wound cellularity was decreased in mice with *S. pneumoniae* infection, which corresponded to loss of neutrophils, monocytes, and macrophages (Figures S6b, S6c and S6d). Wound fluid cytokine and chemokine levels were also reduced in mice with pulmonary *S. pneumoniae* infection (Figure S6e and S6f). Conversely, pulmonary resistance to *S. pneumoniae* infection was not affected by the presence of a wound (Figure S6g). These results indicate that the prioritization of innate immune responses between wound healing and pulmonary infection is not pathogen specific.

### Wounds and lung infection together induce a unique systemic inflammatory response

The bone marrow and circulation are the primary source of monocytes and neutrophils involved in both wound healing and pulmonary antibacterial defense. With competing inflammatory sites, it is possible that there is a limited supply of leukocytes in the bone marrow and/or blood. Therefore, one potential explanation for the decreased cellularity observed in the wounds of wound + *K. oxytoca* mice is that, because large numbers of leukocytes are allocated to the lung, there are fewer available cells to respond to the wound. The number of cells recovered from the bone marrow of wounded, infected, or wound + *K. oxytoca* mice was similar among the three experimental groups with the exception of 48 hours post-infection, in which *K. oxytoca*-infected mice had fewer bone marrow cells, perhaps due to increased mobilization (Figure 3a). Assessing leukocyte populations by flow cytometry analysis, wound and wound + *K. oxytoca* mice had more neutrophils in the bone marrow than infected mice alone at 24- and 48-hours post-infection. The other populations examined were similar among the three experimental groups (Figure 3b and Figure 3c).

**Fig. 3.**
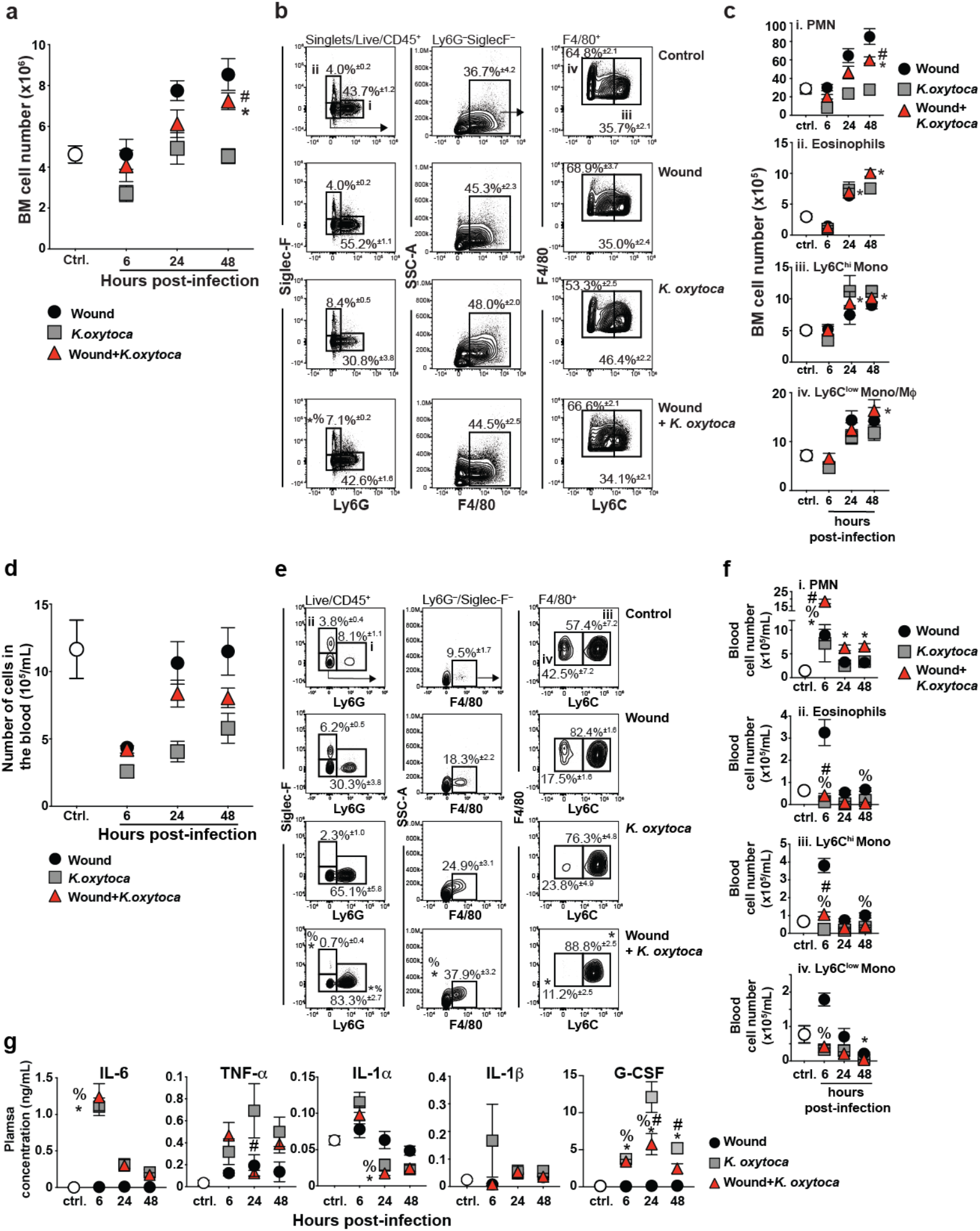
*K. oxytoca* infection alters systemic innate leukocyte and cytokine responses in wounded mice. C57BL/6J mice were wounded by PVA sponge implantation (wound) and a cohort was infected with *K. oxytoca* (wound + *K. oxytoca*) five days later. Additional unwounded mice remained uninfected (control) or were infected intranasally with *K. oxytoca* (*K. oxytoca*). **a)** A time course of showing the total number of cells in the bone marrow (BM). **b)** The proportion of bone marrow Ly6G^+^ neutrophils (PMN, i), Siglec-F^+^ eosinophils (ii), F4/80^+^Ly6C^hi^ monocytes (mono, iii), and F4/80^+^Ly6C^low^ monocytes/macrophages (mono/mΦ) (iv) was assessed by flow cytometry. Representative analyses from 48h post-infection are shown. **c)** The absolute number of bone marrow leukocytes over time. **d)** A time course of the number of cells per milliliter of blood. **e)** The frequency of blood Ly6G^+^ neutrophils (PMN, i), Siglec-F^+^ eosinophils (ii), F4/80^+^ monocytes, F4/80^+^Ly6C^hi^ inflammatory monocytes (mono, iii), and F4/80^+^Ly6C^low^ patrolling monocytes (mono, iv), was determined by flow cytometry analysis. Representative gating from 48h post-infection is shown. **f)** The absolute number of blood leukocytes was assessed over time. **g)** A time course of plasma cytokines was determined by LegendPlex assay (IL-6, TNF-α, IL-1α, and IL-1β) or ELISA (G-CSF). Data are shown as the mean±SEM with minimum n=10 mice per group from three independent experiments. Results are considered statistically significant when p ≤ 0.05. Statistically significant changes between control and wound + *K. oxytoca* are denoted by *, between wound and wound + *K. oxytoca* are denoted by %, and between *K. oxytoca* and wound + *K. oxytoca* are denoted by #.

Examining the blood, all three experimental groups had a decrease in the number of circulating cells compared to control mice, likely due to margination to the inflamed peripheral sites (Figure 3d). Flow cytometry analysis revealed that wound + *K. oxytoca* mice had significantly more circulating neutrophils 6 hours post-infection compared to wounded or infected mice alone, and their levels remained elevated 24- and 48-hours post-infection (Figures 3e, 3f, and S7). In contrast, circulating eosinophils as well as Ly6C^hi^ and Ly6C^low^ monocytes were decreased in wound + *K. oxytoca* mice compared to wounded mice alone (Figures 3e and 3f, and Figure S7). This suggests that, in wound + *K. oxytoca* mice, a limiting systemic supply of Ly6C^hi^ monocytes may contribute to their decline in the wound, while the abundance of circulating neutrophils drives their rapid accumulation in the BALF.

Systemic cytokine levels are important in the activation of immune cells. To assess whether competing insults altered the balance of systemic cytokines, the plasma concentration of inflammatory cytokines was measured by ELISA or multiplex bead assay. Mice were wounded by PVA sponge implantation and/or infected as previously described (Figure 1c). None of the cytokines examined were induced in control or wounded mice at the time points examined. IL-6 was strongly induced 6 hours after *K. oxytoca* infection with no effect of prior wounding. TNF-α was also elevated systemically in mice infected with *K. oxytoca*, but the induction of TNF-α was transiently suppressed in wound + *K. oxytoca* mice at 24h post-infection. The concentration of IL-1α and IL-1β in the plasma of wound + *K. oxytoca* mice followed the pattern observed in infected mice alone (Figure 3g). G-CSF was rapidly induced in the plasma of all infected groups as early as 6 hours post-infection. This could explain the increase in blood neutrophil content in wound + *K. oxytoca* mice, as these mice have both increased bone marrow neutrophil content driven by the wound response and an infection-induced rise in G-CSF, which regulates neutrophil mobilization (*39*–*41*). Taken together, these data indicate that a deficit in the number of circulating monocytes contributes to the loss of monocyte cellularity in the wound; in contrast, the abundance of circulating neutrophils is not consistent with their absence in the wound environment, suggesting other factors contribute to this phenotype.

### Fewer leukocytes migrate to the wounds of mice with pulmonary infection

The loss of wound cellularity in wound + *K. oxytoca* mice could stem from a decreased ability of circulating leukocytes to migrate to the wound site. To test this, a bone marrow cell adoptive transfer approach was taken. CD45.2^+^ C57BL/6J recipient mice were wounded by PVA sponge implantation, and a cohort was infected intranasally with *K. oxytoca* 5 days later. Twenty-four hours after infection, bone marrow cells isolated from naive CD45.1 congenic mice were transferred intravenously to wound or wound + *K. oxytoca* CD45.2^+^ recipient mice (Figure 4a). The fraction of CD45.1^+^ donor-derived cells in the wounds of recipient mice was assessed 48 hours post-infection by flow cytometry analysis. There were fewer CD45.1^+^ donor-derived cells by proportion and total number in the wounds of wound + *K. oxytoca* mice as compared to the wounds of wounded recipient mice alone (Figure 4b). Similarly, there were significantly fewer donor derived CD45.1^+^ Ly6G^+^ neutrophils and F4/80^+^Ly6C^hi^ monocytes by proportion and absolute number in the wounds of wound + *K. oxytoca* recipient mice compared to wounded recipient mice alone (Figure 4c and 4d). There was only a very small proportion (<0.05%) of CD45.1^+^F4/80^+^Ly6C^low^ macrophages in recipient wounds (Figure 4d); these were likely derived from monocytes that matured *in situ* after migrating from the circulation (*20*). Together, these data indicate that circulating neutrophils and inflammatory monocytes are impaired in their ability to migrate to wounds in mice with an ongoing *K. oxytoca* infection.

**Fig. 4.**
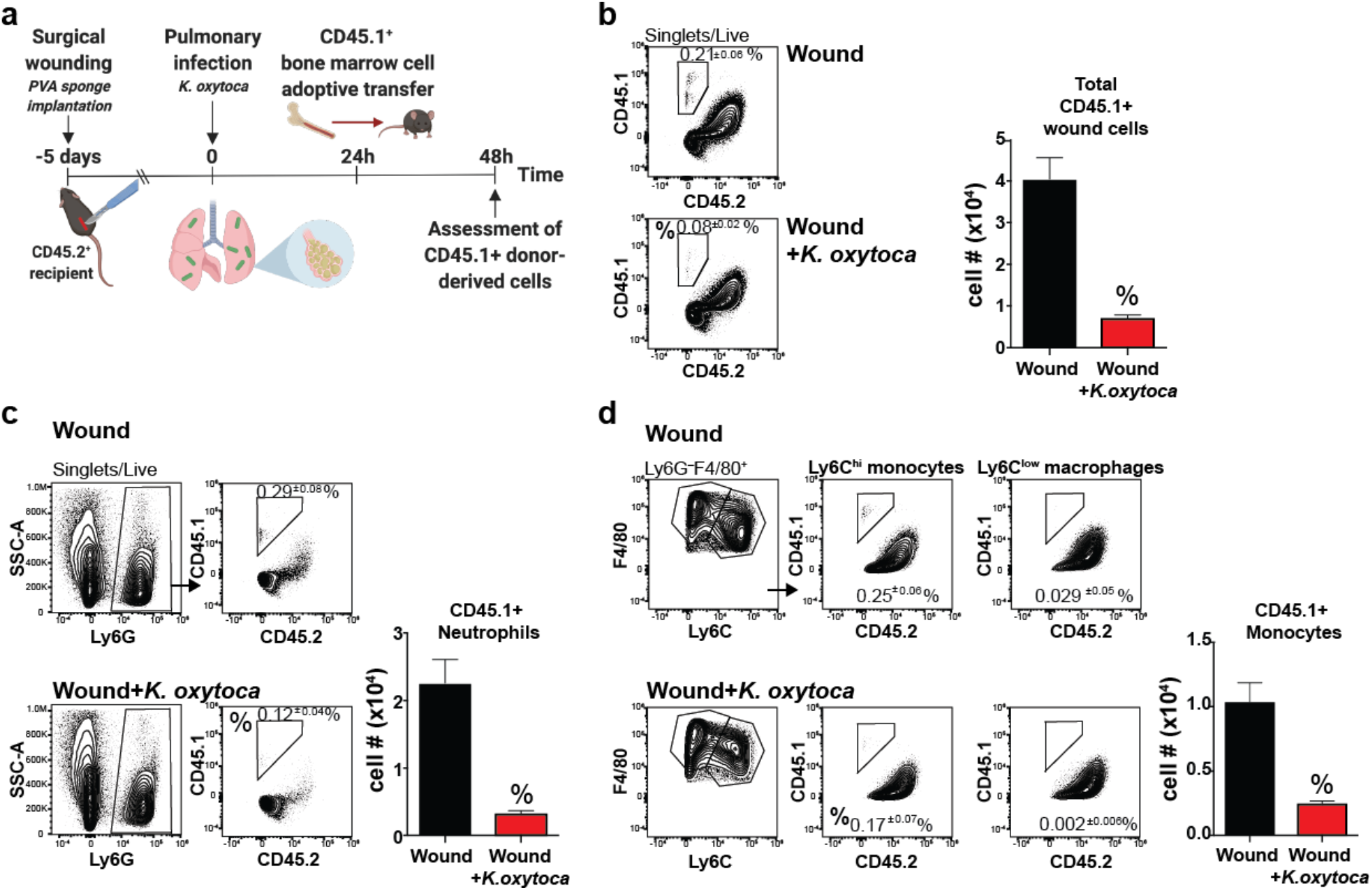
Neutrophil and monocyte trafficking to wounds is impaired in mice with pulmonary *K. oxytoca* infection. **a)** Naïve CD45.1^+^ congenic bone marrow cells were adoptively transferred to uninfected or *K. oxytoca*-infected mice with PVA sponge wounds. **b)** The percentage and number of donor-derived CD45.1^+^ cells in the wounds of uninfected and *K. oxytoca*-infected mice, as determined by flow cytometry. **c)** The percentage and number of donor-derived CD45.1^+^ Ly6G^+^ neutrophils in the wounds of uninfected and *K. oxytoca*-infected mice, as determined by flow cytometry. **d)** The percentage and number of donor-derived CD45.1^+^ F4/80^+^Ly6C^hi^ monocytes and the percentage of CD45.1^+^ F4/80^+^Ly6C^low^ macrophages in the wounds of uninfected and *K. oxytoca*-infected mice, as determined by flow cytometry. The number of macrophages is excluded due to low frequency. Data are shown as the mean±SEM with minimum n=12 mice per group from three independent experiments. Results are considered statistically significant when p ≤ 0.05. Statistically significant changes between wound and wound + *K. oxytoca* are denoted by %.

### Pulmonary infection rapidly suppresses wound chemokine signals

The decrease in the accumulation of adoptively transferred cells to the wounds of infected mice could be due to a local or systemic imbalance of chemokine signals. To assess this, we examined the expression of chemokines and chemokine receptors that are important in neutrophil and monocyte migration to inflamed sites. Mice were wounded and/or infected as previously described (Figure 1c). The neutrophil chemoattractants CXCL1 and CXCL5 were measured in the wound fluid, BALF, and plasma of all experimental groups. In the wound fluid, CXCL1 and CXCL5 were rapidly suppressed within 6 hours of *K. oxytoca* infection and never recovered to the levels measured in wounded mice alone at the time points examined (Figure 5a). CXCL1 was induced in the BALF and the plasma in response to *K. oxytoca* infection. Interestingly, CXCL1 concentrations were suppressed in both of these compartments in wound + *K. oxytoca* mice compared to infected mice alone at one or more time points. While CXCL5 was high in the plasma of all experimental groups, it was induced only by infection in the BALF (Figure 5b and c).

**Fig. 5.**
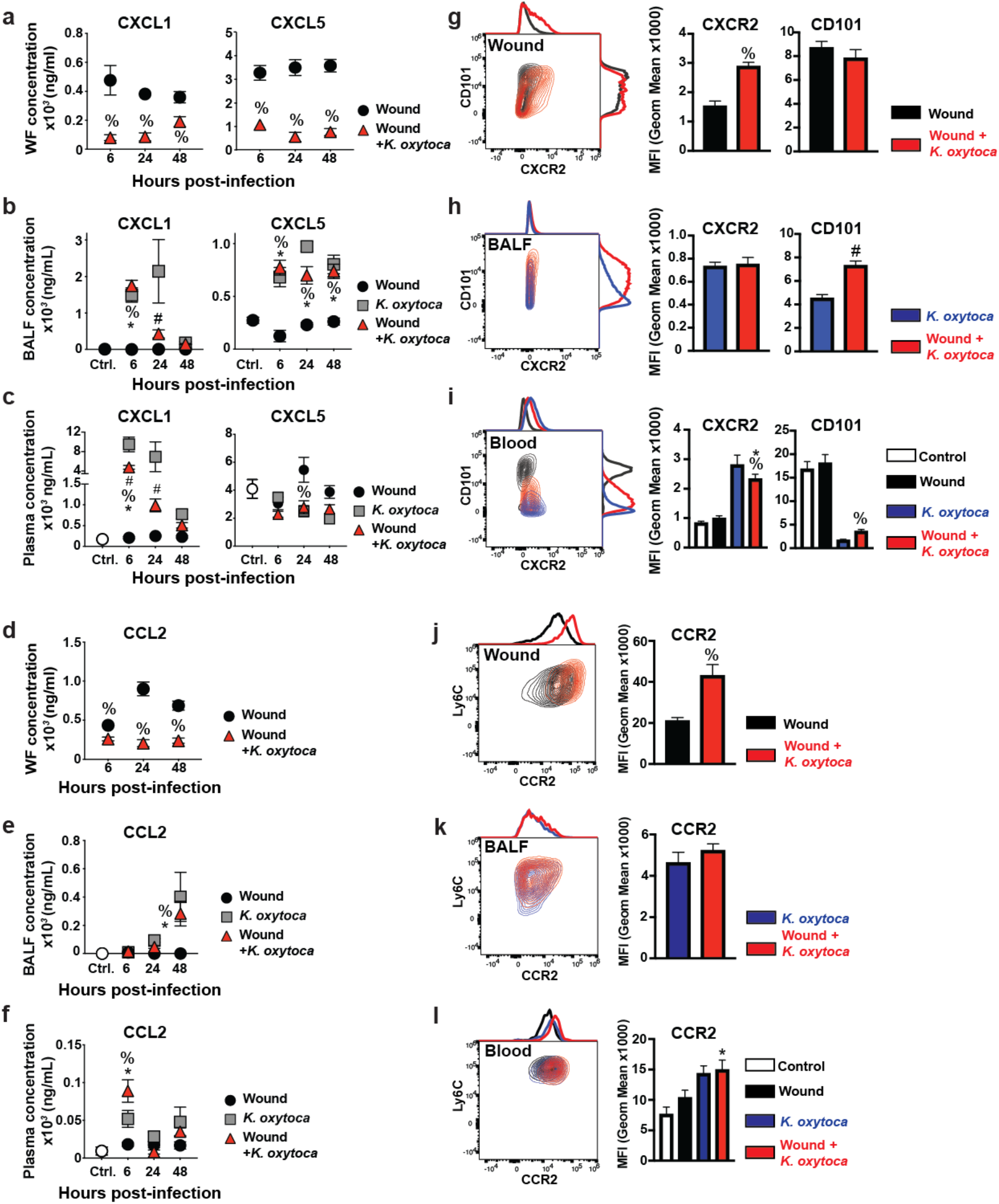
Wounds and pulmonary infection disrupt local and systemic chemokine responses. C57BL/6J mice were wounded by PVA sponge implantation (wound) and a cohort was infected 5 days later with *K. oxytoca* (wound + *K. oxytoca*). Additional unwounded mice remained uninfected (ctrl.) or were infected with *K. oxytoca* (*K. oxytoca*). **a-c)** A time course of CXCL1 and CXCL5 was determined in the wound fluid (WF) (a), BALF (b), and plasma (c) by ELISA. **d-f)** A time course of CCL2 was determined in the wound fluid (d), BALF (e), and plasma (f) by LegendPlex assay. **g-i)** Expression of CD101 and CXCR2 was determined on neutrophils isolated from the wound (g), BALF (h), and blood (i) by flow cytometry analysis. **j-l)** Expression of CCR2 was determined on monocytes isolated from the wound (i), BALF (k), and blood (l) by flow cytometry analysis. Data are shown as the mean±SEM with a minimum n=7 mice per group from at least two independent experiments. Results are considered statistically significant when p ≤ 0.05. Statistically significant changes between control and wound + *K. oxytoca* are denoted by *, between wound and wound + *K. oxytoca* are denoted by %, and between *K. oxytoca* and wound + *K. oxytoca* are denoted by #.

A similar rapid and sustained reduction of the monocyte chemoattractant CCL2 was observed in the wound fluid within 6 hours post-infection (Figure 5d). As with CXCL1 and CXCL5, this suppression of wound fluid CCL2 expression preceded the reduction in wound cellularity, which began at 24 hours post-infection (Figure 1). In the BALF and the plasma, CCL2 levels were highest in infected groups, peaking at 48- and 6-hours post-infection, respectively (Figure 5e and f).

Inflammation can alter the phenotype of innate leukocytes, which may have functional consequences at the tissue level. In the steady state, neutrophils in the circulation are phenotypically mature and express high levels of the marker CD101, while inflammation can lead to the release of CD101^low^ neutrophils from the bone marrow that are immature and functionally distinct (*40*). Mice were wounded and infected as described above (Figure 1c), and neutrophils from the wound, BALF, and blood were examined for expression of CD101 expression and CXCR2, the receptor for CXCL1 and CXCL5 (*42*). The neutrophils that were recruited to the wounds of wound + *K. oxytoca* mice had significantly higher expression of CXCR2, while their baseline high CD101 status was not altered, compared to wounded mice alone (Figure 5g). In contrast, CXCR2 expression on BALF neutrophils was not impacted by the presence of a wound; however, the presence of a wound led to slightly increased expression of CD101 (Figure 5h). To understand whether these changes originated from systemic effects, blood neutrophils were also examined. Blood neutrophils isolated from wounded mice were primarily CXCR2^low^CD101^hi^. In contrast, in infected mice, blood neutrophils had higher CXCR2 and lower CD101 expression. Blood neutrophils from wound + *K. oxytoca* mice resembled those from infected mice alone, although expression of both markers trended toward an intermediate phenotype (Figure 5i). Thus, infection drove the increased CXCR2 expression seen in the wounds of infected mice, while the presence of a wound caused slightly elevated CD101 expression on blood neutrophils, which was also evident in the BALF.

Monocytes were similarly examined for changes in the expression of CCR2, the receptor for CCL2, to determine whether this could contribute to their impaired migration to the wounds of infected mice. Inflammatory monocytes isolated from the wounds of infected mice demonstrated a greater than two-fold increase in their expression of CCR2 compared to wounded mice alone (Figure 5j). In contrast, in the BALF of infected mice, CCR2 expression on monocytes was not affected by prior wounding (Figure 5k). Systemically, infection led to an increase in CCR2 expression on circulating monocytes (Figure 5l), which was reflected in the wound monocyte compartment in wound + *K. oxytoca* mice. Interestingly, in this group, the CCR2 MFI of wound monocytes was nearly three-fold higher than that measured on blood monocytes. This trend was also observed in neutrophil CXCR2 expression, albeit it to a lesser degree. These data suggest that leukocytes with higher chemokine receptor expression are selectively recruited to the chemokine-poor environment, or that a lack of negative feedback prevents their downregulation. Taken together, these data demonstrate that pulmonary infection and wound healing responses drive unique systemic changes in the expression of chemokines and their receptors, which have consequences at the tissue level.

### Restoring innate immune cell trafficking to wounds with exogenous CCL2 and CXCL1 rescues wound healing

Given the reduction in monocyte and neutrophil trafficking to the wounds of wound + *K. oxytoca* mice, we hypothesized that restoring trafficking would improve wound healing responses in these mice. To examine the effect of CCL2 and CXCL1 administration on wound innate leukocyte responses, the PVA sponge implantation model was used. Mice were wounded and/or infected as described above (Figure 1c). Recombinant CCL2 and recombinant CXCL1 were mixed and injected into each implanted sponge of wound + *K. oxytoca* mice at the time of infection and again 24 hours post-infection. An additional cohort of wound + *K. oxytoca* mice, as well as uninfected mice, received PBS vehicle injections in the implanted sponges. CCL2 and CXCL1 treatment increased the wound cellularity of wound + *K. oxytoca* mice at 48 hours post-infection, compared to wound + *K. oxytoca* mice treated with PBS (Figure 6a). The frequency and number of neutrophils was significantly increased in wound + *K. oxytoca* mice with the addition of exogenous chemokines, while the number of monocytes was not significantly influenced despite the addition of the monocyte chemoattractant CCL2 (Figure 6b and 6c). Wound chemokine treatments did not affect the number of circulating, lung tissue, or BALF cells, including neutrophils and monocytes (Figure 6d-i). Bacterial titers were elevated in the infected mice that received wound chemokine treatments (Figure 6j), indicating that the redirection of neutrophil trafficking towards the wound had a detrimental effect on pulmonary bacterial resistance.

**Fig. 6.**
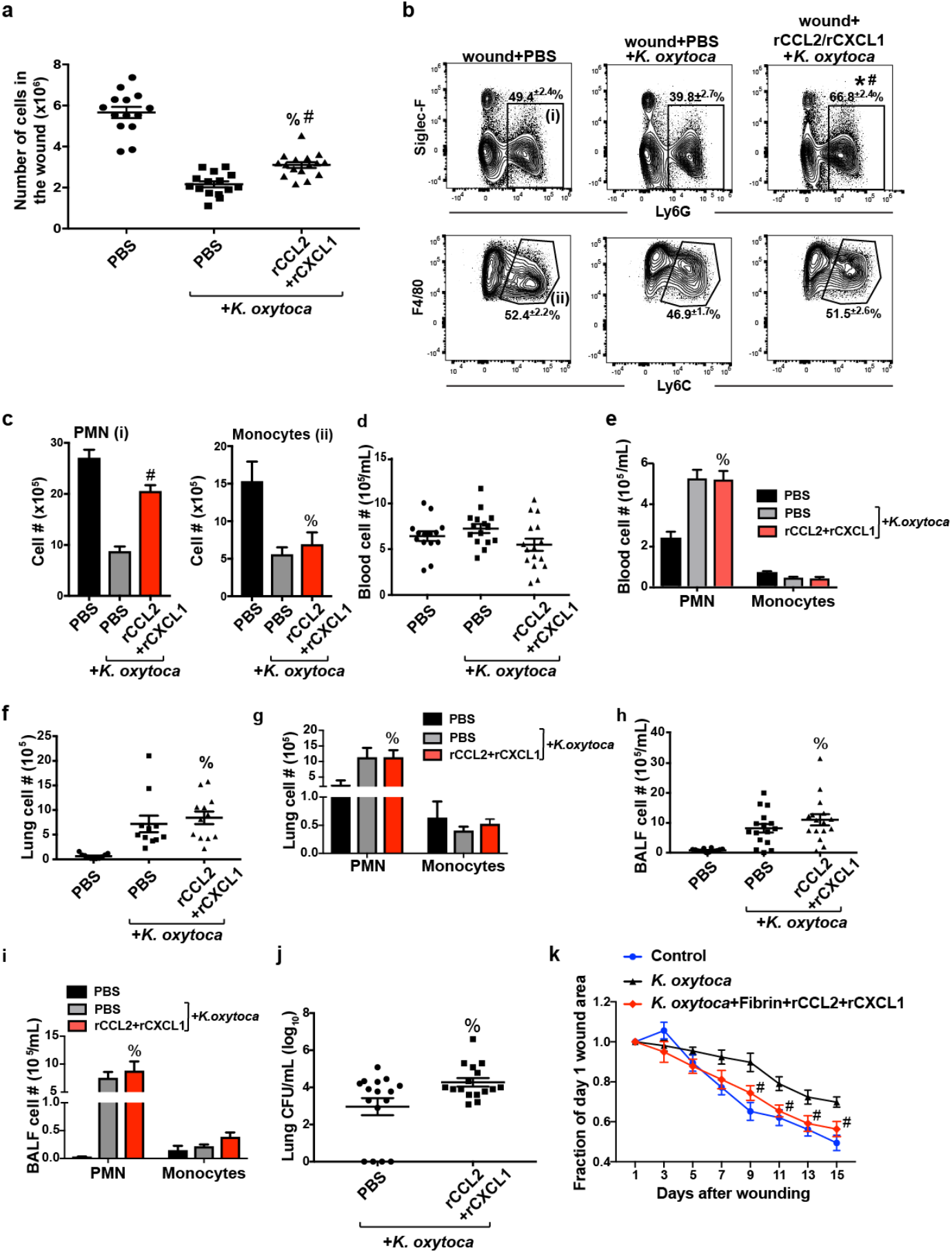
Addition of exogenous CCL2 and CXCL1 to wounds improves healing at the expense of pulmonary resistance to *K. oxytoca* infection. Mice with PVA sponge wounds were uninfected or infected with *K. oxytoca* five days later. Sponges in infected mice were injected with PBS vehicle or recombinant CCL2+CXCL1 at the time of infection and 24h post-infection. **a)** Wound and blood cellularity was determined 48h post-infection. **b-c)** The effect of chemokine administration on the percentage (b) and absolute number (c) of wound Ly6G^+^ neutrophils and F4/80^+^Ly6C^hi^ monocytes was determined by flow cytometry analysis at 48h post-infection. **d-e)** The effect of wound chemokine treatments on blood cellularity is shown in (d) and the number of blood neutrophils and inflammatory monocytes is shown in (e) at 48h post-infection. **f-g)** The cellularity of the lung tissue (f), the number of lung Ly6G^+^ neutrophils and F4/80^+^Ly6C^hi^ monocytes (g) was determined at 48h post-infection. **h-i)** The cellularity of the BALF (h) and number of BALF Ly6G^+^ neutrophils and F4/80^+^Ly6C^hi^ monocytes (i) was determined in mice at 48h post-infection. **j)** The effect of wound chemokine treatments on lung bacterial burden at 48h post-infection was determined by CFU analysis. **k)** Excisional tail wounds were performed on C57BL/6J mice. One cohort of wounded mice remained uninfected (control) and a cohort was infected with *K. oxytoca*. A mixture of recombinant CCL2 and CXCL1 was applied to the wound beds of a subset of *K. oxytoca-*infected mice using a fibrin vehicle (*K. oxytoca*+Fibrin+rCCL2+rCXCL1). The area of the tail wound was measured to determine the effect of chemokine application on the rate of wound closure. Data are shown as the mean±SEM with minimum n=12 mice per group from three independent experiments. Results are considered statistically significant when p ≤ 0.05. In (a-j), % indicates a statistically significant change between wound + PBS and wound + rCCL2/rCXCL2 + *K. oxytoca*, and # indicates a statistically significant change between wound + PBS + *K. oxytoca* and wound + rCCL2/rCXCL2 + *K. oxytoca.* In (k), # denotes a statistically significant change between *K. oxytoca* and rCCL2/rCXCL2 + *K. oxytoca*.

To determine whether the addition of exogenous chemokines could improve the rate of wound healing, recombinant CCL2 and CXCL1 were delivered to the tail wounds of *K. oxytoca*-infected mice using a topical fibrin sealant. Recombinant CCL2 and CXCL1 were mixed and incorporated into the fibrin sealant prior to application. Fibrinolysis of the sealant by wound site proteases delivers the incorporated chemokines to the wound bed. The wounds of uninfected mice were left untreated (control) or were treated with fibrin sealant. A subset of tail-wounded mice was infected with *K. oxytoca* on wound day 1. The wound beds of tail wound + *K. oxytoca* mice were untreated, treated with fibrin sealant, or treated with fibrin sealant containing recombinant CCL2 and recombinant CXCL1. Treatments were applied every day from wound days 1 to 7, then every other day from days 9 to 15. *K. oxytoca*-infected mice had the slowest healing tail skin wounds (Figure 6k). Application of fibrin sealant to the tail skin wounds of control and *K. oxytoca*-infected mice did not significantly affect the rate of healing (Figure S8). However, treatment of tail skin wounds with fibrin supplemented with recombinant CCL2 and recombinant CXCL1 restored the rate of healing in *K. oxytoca*-infected mice to that of the control group (Figure 6k). Together, these data indicate that chemokine-mediated signals regulate innate leukocyte recruitment to the site of wounding and the rate of tail skin wound closure.

## Discussion

This work investigated the concept of innate immune prioritization of inflammatory sites, which is essential in the full understanding of the innate immune response given its multiple roles in health and disease. Building upon previous work, which demonstrated that pulmonary infection with influenza A virus in mice suppresses wound healing in the skin (*30*), the current work identifies the mechanisms underlying impaired wound healing in mice with bacterial pulmonary infection. We show that post-operative human patients with pneumonia had decreased wound healing, as indicated by an increased rate of dehiscence. Innate leukocytes are critical to the repair of injured tissue and to the early control of pulmonary infection, so to investigate the role played by the innate immune system, we developed a murine model of post-operative pulmonary infection. In this model, the innate immune response is faced with two distal and competing inflammatory insults: one in the skin and one in the lung. There is considerable overlap in the cellular and cytokine responses that orchestrate acute wound healing and the pulmonary response to bacterial infection (*11*–*13*, *15*–*17*, *20*, *21*, *28*, *36*, *43*–*51*); therefore, we hypothesized that one inflammatory site would more strongly recruit innate leukocytes, and thus take priority over the other in a concept we call “innate immune triage.”

It has been shown in other systems that disruption of innate immune cellular responses can alter or delay wound healing (*9*, *12*, *13*, *15*, *16*, *18*, *43*). In murine models, as we observed in our patient data, the pulmonary response to bacterial infection was prioritized at the expense of cutaneous wound healing. Our data demonstrated that the healing rate of excisional tail wounds was delayed following the onset of pulmonary *K. oxytoca* infection. Examining the cellular mechanisms of impaired healing, we determined that chemokine levels in PVA sponge wounds of infected mice dropped as early as 6 hours post-infection, before any observed decline in wound cellularity. This suggests that wound chemokines are actively suppressed by signals from the infected lung. This dip in chemokine levels was followed by greatly reduced wound cellularity and cytokine concentrations. The loss of wound cellularity was attributed primarily to a decrease in neutrophils and inflammatory monocytes. These leukocytes migrate from the blood to the wound site, where they clear injured tissue debris, coordinate inflammatory responses, and, in the case of monocytes, differentiate into wound macrophages to drive repair responses (*12*, *15*, *20*). Adoptive transfer experiments demonstrated that a decrease in leukocyte migration contributed to the loss of monocytes and neutrophils in the wounds of infected mice. These findings are consistent with studies that show blocking the early acute cellular responses to wounding disrupts the later stages of healing (*9*, *13*, *14*, *17*, *43*).

It has been reported that injury or infection can induce systemic immunosuppression, which we hypothesized could contribute to the impaired wound healing that occurred in mice with pulmonary bacterial infection (*26*, *27*, *52*, *53*). Evidence of this was seen in the delayed expression of TNF-α and CXCL1 in the plasma of wound + *K. oxytoca* mice compared to infected mice alone. TNF-α has coordinated expression with many chemokines, including CXCL1, suggesting that the deficit in these factors may be linked (*54*–*57*). A similar trend in TNF-α and CXCL1 induction was observed in the BALF, indicating that the systemic effect was driven by a transient suppression of local lung cytokine and chemokine signaling. Surprisingly, the suppressive effect of wounding on BALF cytokine and chemokine levels during *K. oxytoca* infection did not have an overtly detrimental effect on BALF cellularity or the control of bacterial infection. Perhaps this is due to high vascularization in the lung (*58*), permitting even low levels of chemokines to attract an adequate number of cells to respond to infection. This is likely why wound + *K. oxytoca* mice were able to mediate early control over *K. oxytoca* infection at the time points examined. These results are in contrast to what has been reported in cases of severe trauma, which can cause immune dysfunction and impaired pulmonary immune responses, thereby increasing the risk of developing secondary lung infection (*2*, *27*, *59*). The wound models implemented in this study do not generate a strong or sustained systemic inflammatory response and do not recapitulate trauma-induced immune suppression, which is consistent with the lack of effect on the pulmonary antibacterial response in wound + *K. oxytoca* mice.

Despite the transient depression in systemic cytokine and chemokine signaling, wounding bolstered blood cellularity in both uninfected and infected mice. In particular, there were twice as many circulating neutrophils in wound + *K. oxytoca* mice compared to other treatment groups. Despite the surplus of circulating neutrophils, the wounds of wound + *K. oxytoca* mice had very few neutrophils compared to the wounds of uninfected mice. This indicates that a lack of neutrophil chemotactic signal from the wound site was responsible for the decreased number of wound neutrophils in wound + *K. oxytoca* mice. In contrast, the number of circulating Ly6C^hi^ monocytes was lowest in wound + *K. oxytoca* mice, so the decrease in wound monocyte number in *K. oxytoca*-infected mice may reflect both a decrease in the circulating supply and a loss of chemotactic signal from the wound. Furthermore, it was found that mice with competing inflammation in the skin and lung had a unique phenotype regarding the distribution of cells in the bone marrow and blood, as well as the activation state of these cells, which may also contribute to functional changes in the tissue.

We hypothesized that redirecting neutrophil and monocyte trafficking to the wounds of wound + *K. oxytoca* mice would improve healing. Serial application of recombinant CCL2 and CXCL1 to excisional tail skin wounds accelerated the rate of wound closure in *K. oxytoca*-infected mice to that of uninfected mice. Injection of recombinant CCL2 and CXCL1 into PVA sponge wounds improved wound cellularity in wound + *K. oxytoca* mice through an increase in neutrophils. Neutrophils were the predominant circulating leukocyte population in infected mice, and this is likely why they were preferentially recruited to chemokine-treated wounds. Interestingly, wound + *K. oxytoca* mice that received wound chemokine treatments showed a slight impairment in bacterial clearance in the lungs, indicating that the redistribution of neutrophils to the wound altered pulmonary resistance to bacterial infection. Surprisingly there was not a decrease in the overall cellularity of the lung or BALF, including neutrophils and monocytes; the reason for this loss of resistance to bacterial infection will require further investigation but was perhaps driven by impaired activation or bactericidal activity of the cells. Given the growing appreciation of neutrophil heterogeneity linked to function, the finding that neutrophils in wound + *K. oxytoca* mice display altered phenotypes systemically and locally support this concept (*60*).

This work provides insight into how the innate immune response is equipped to handle simultaneous distal inflammatory insults. Clinical data suggested that surgical patients who acquire pneumonia do not heal as well. This is important because delayed wound healing leaves patients susceptible to a variety of complications, including wound infection or systemic secondary infections (*2*, *26*), hernias, debilitating scar tissue formation (*61*), permanent disablement, and increased mortality (*62*, *63*). Furthermore, the treatment of poorly healing wounds presents a major economic burden to society and the healthcare system (*64*, *65*). Modeling this situation in mice, we found that the innate immune response can indeed prioritize its response to one inflammatory site over another. Pulmonary infection drove a rapid and dramatic suppression of chemokine signals in the wound, which manifested in a breakdown of early acute wound healing responses mediated by the innate immune response. Additionally, this study demonstrates how competing inflammatory insults drive unique phenotypes in circulating leukocytes, which may be linked to functional deficits in downstream tissue responses. Treating wounds with chemokines improved healing, but this occurred at the expense of bacterial clearance in the lung. These findings demonstrate that distal inflammatory insults compete for innate immune cellular resources, and prioritization of the immune response towards one inflamed site over the other is dictated by chemokine-mediated signals. With the immune response directed towards the lungs, poorly healing wounds in patients with hospital-acquired pneumonia may contribute to their increased morbidity. This study introduces the potential of using chemokine-based treatments to manipulate the prioritization of innate leukocyte responses to improve wound healing in high-risk patient populations. Overall, this work provides a mechanistic understanding of innate immune function in complex inflammatory contexts, which has broad implications in cases of clinical comorbidities and in understanding immune responses in a systemic context.

## Acknowledgments

The authors would like to thank Christine Biron and Kayla Campbell for helpful discussions, commentary, and editorial assistance on the manuscript, as well as Kevin Carlson and the Brown University Flow Cytometry and Sorting Facility for valuable flow cytometry consultation and assistance.

## Author contributions

M.J.C. designed the experiments, performed most of the experiments, and wrote the paper, Y.X. designed the experiments and performed most of the experiments, S.F.M. and B.M.H. obtained and analyzed the NSQIP data, J.E.A. designed the experiments, W.L.H. performed the mouse surgeries and assisted in other experiments, H.L.T., K.R.P.C, A.R.D.J, and L.C. assisted with experiments, A.M.J designed the study, designed the experiments, and wrote the paper.

## Data and materials availability

All relevant data is included in the manuscript and supplementary data. Materials will be made available upon request.

