## Supplemental Files for "Pulmonary infection interrupts acute cutaneous wound healing through disruption of chemokine signals"

### Supplementary Data

**Supplementary Table 1. Rate of surgical site infection among surgical patients with or without pneumonia**

| PNEUMONIA | Surgical Site Infection | | TOTAL |
|---|---|---|---|
| | No Infection | Infection | |
| No Pneumonia | 80464 (93.7%) | 5413 (6.30%) | 85877 |
| Pneumonia | 3479 (93.2%) | 252 (6.75%) | 3731 |

**Supplementary Table 2. Rate of abdominal wound dehiscence among surgical patients with urinary tract infection**

| Urinary Tract Infection (UTI) | DEHISCENCE | | TOTAL |
|---|---|---|---|
| | No Dehiscence (% of Total) | Dehiscence (% of Total) | |
| No UTI | 86102 (98.60%)^ | 1220 (1.40%)^ | 87322 |
| UTI | 2251 (98.47%)# | 35 (1.53%)# | 2286 |

^Percent of total number of patients in 'No UTI' group
#Percent of total number of patients in 'UTI' group

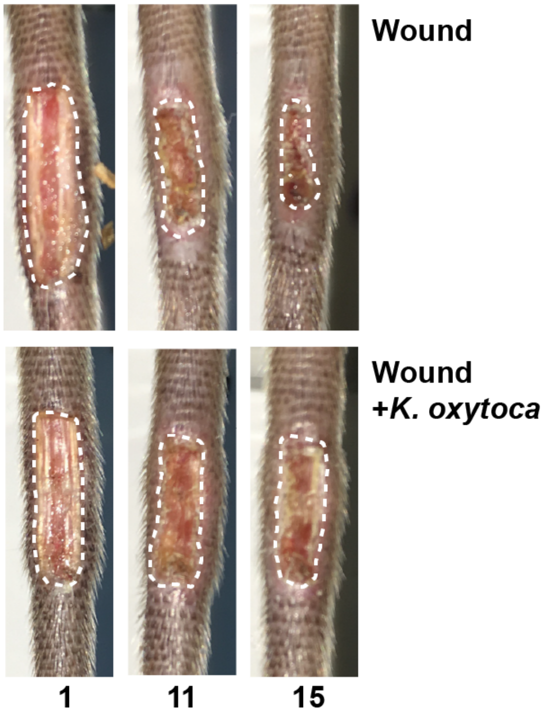


**Figure S1. The effect of pulmonary *K. oxytoca* infection on excisional tail wound closure.** Enlarged representative images of excisional tail wounds from uninfected control mice (top) or mice with pulmonary *K. oxytoca* infection (bottom) on wound days 1, 11, and 15 are shown, with wound margins traced for clarity.

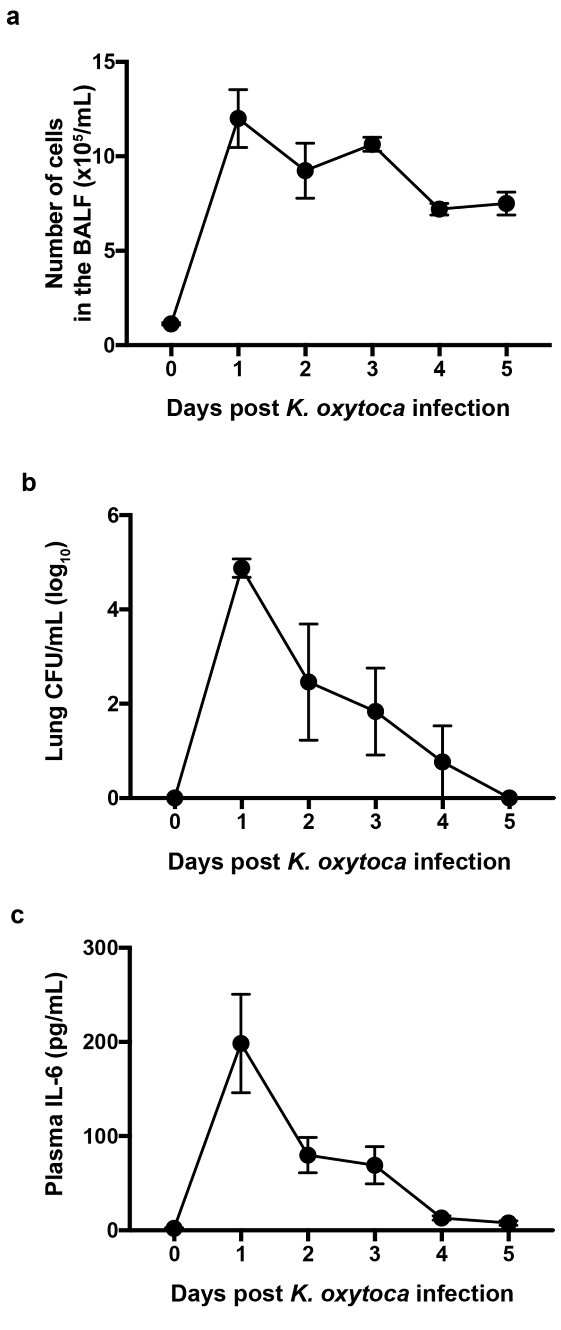


**Figure S2. Time course of pulmonary *K. oxytoca* infection.** The kinetics of BALF cellularity (a), lung bacterial titers (b), and plasma IL-6 concentration (c) demonstrate the course of infection and inflammation in mice infected intranasally with $3x10^7$ CFU *K. oxytoca.* The peak response occurs one day after infection for all parameters. N = 3-7 mice per group.

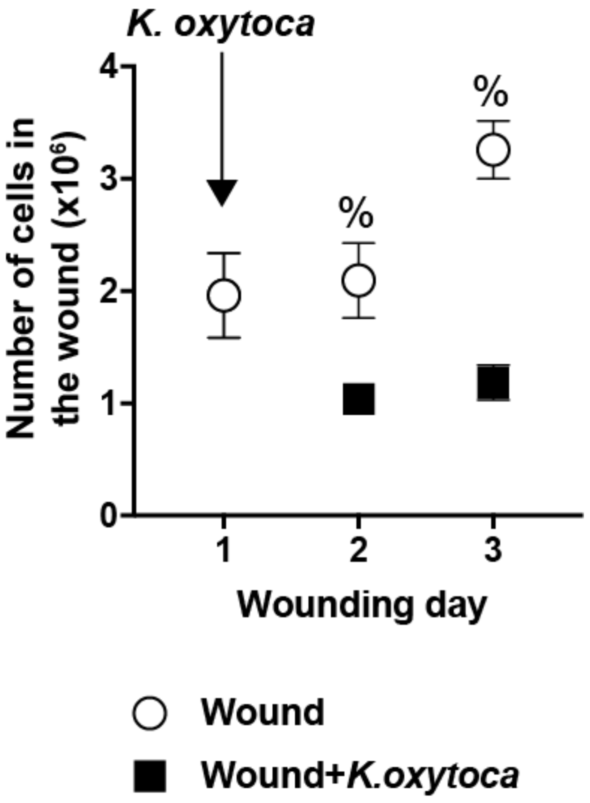


**Figure S3. Pulmonary *K. oxytoca* infection causes reduced wound cellularity at early wound time points.** Pulmonary infection was initiated in a cohort of wounded mice on wound day 1, and wound cell number was recorded on wound days 1, 2, and 3. Data are shown as the mean±SEM with n=12 mice per group from three independent experiment. % indicates a statistically significant change between wound and wound + *K. oxytoca*.

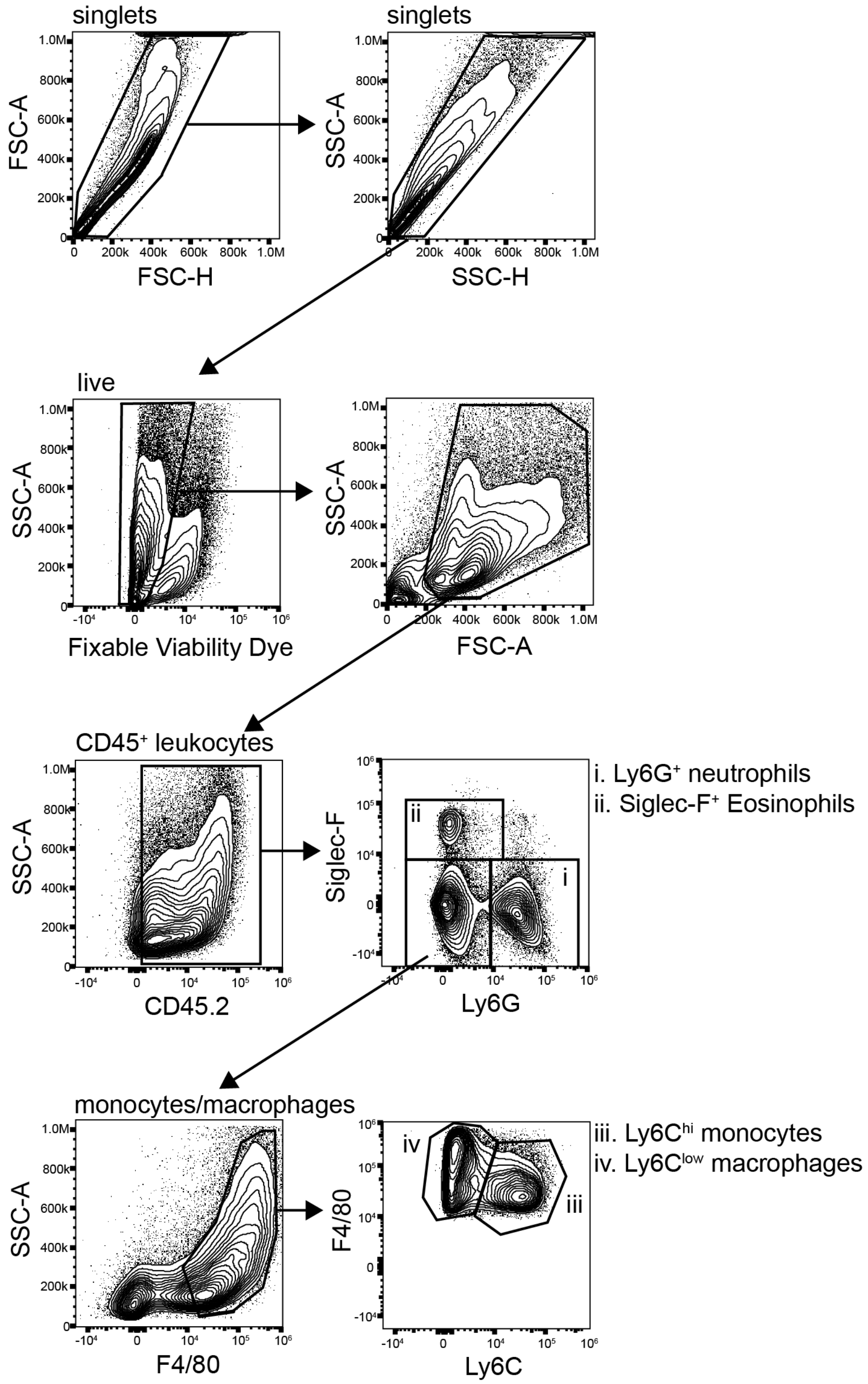


**Figure S4. Representative flow cytometry gating strategy to identify wound innate leukocytes.** The following base gating strategy was employed to quantify innate leukocytes in the wound. Following doublet exclusion, dead cells were removed from the analysis using a fixable viability dye. Cell debris and residual red blood cells were excluded by size using FSC-A and SSC-A parameters. Hematopoietic cells were identified as CD45.2$^+$. Neutrophils (i) were identified as Ly6G$^+$Siglec-F$^-$. Eosinophils (ii) were identified as Siglec-F$^+$Ly6G$^-$. F4/80$^+$ monocytes/macrophages were gated from the Ly6G$^-$Siglec-F$^-$ population. F4/80$^+$ cells were fractionated into Ly6C$^{hi}$ monocytes (iii) and Ly6C$^{low}$ macrophages (iv). A day 7 wound is shown.

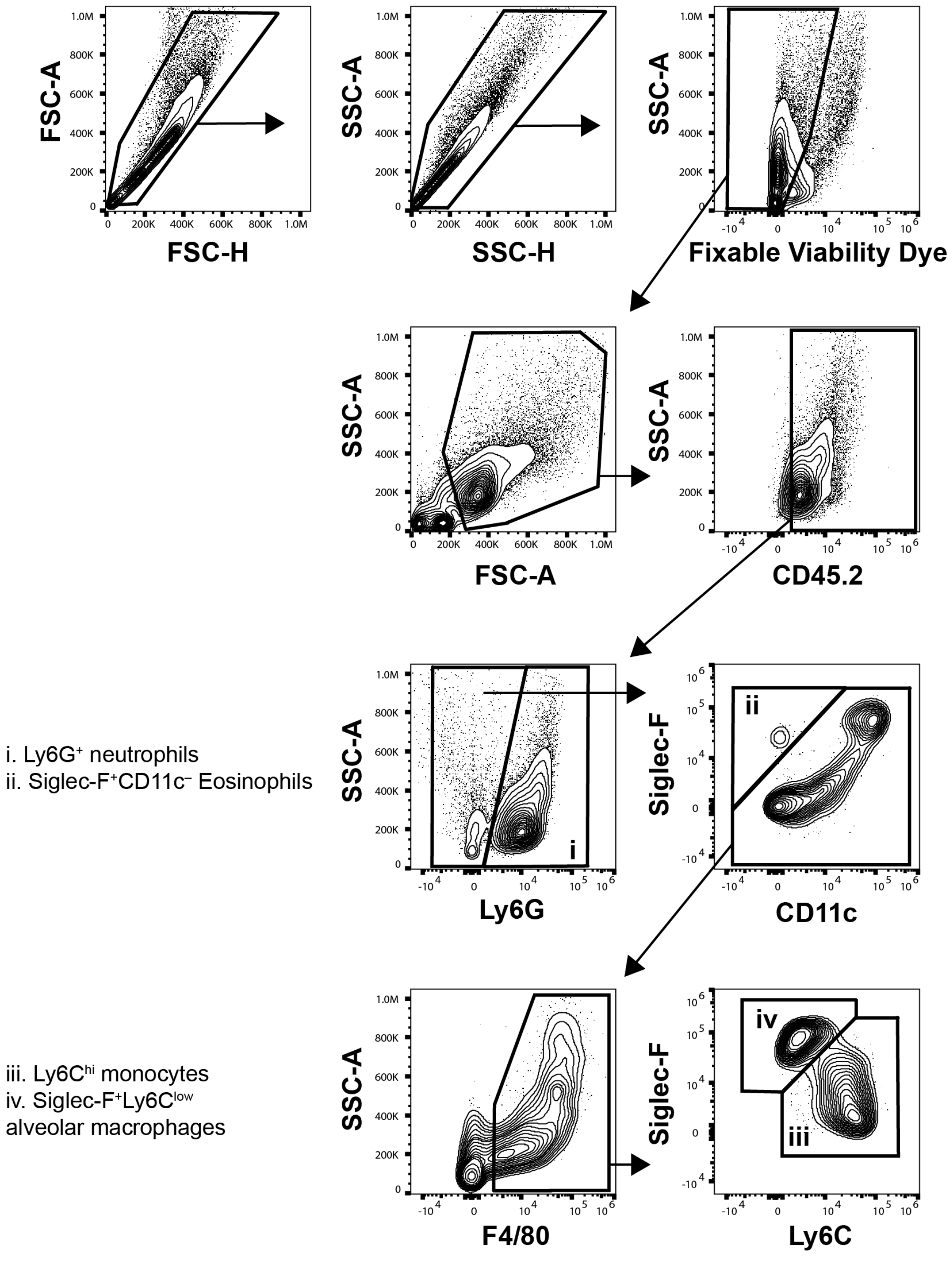


**Figure S5. Representative flow cytometry gating strategy to identify BALF innate leukocyte populations.** This gating strategy was employed to quantify innate leukocytes in the BALF. Doublets were excluded, then dead cells were removed from the analysis using a fixable viability dye. Cell debris and residual red blood cells were excluded by size using the FSC-A and SSC-A parameters. Hematopoietic cells were identified as CD45.2$^+$. Neutrophils (i) were identified as Ly6G$^+$. Eosinophils (ii) were identified as Siglec-F$^+$CD11c$^-$Ly6G$^-$. F4/80$^+$ monocytes/macrophages were gated from the Ly6G$^-$Siglec-F$^-$ population. F4/80$^+$ cells were fractionated into Ly6C$^{hi}$ monocytes (iii) and Siglec-F$^+$Ly6C$^{low}$ alveolar macrophages (iv). BALF cells from 48h post-infection is shown.

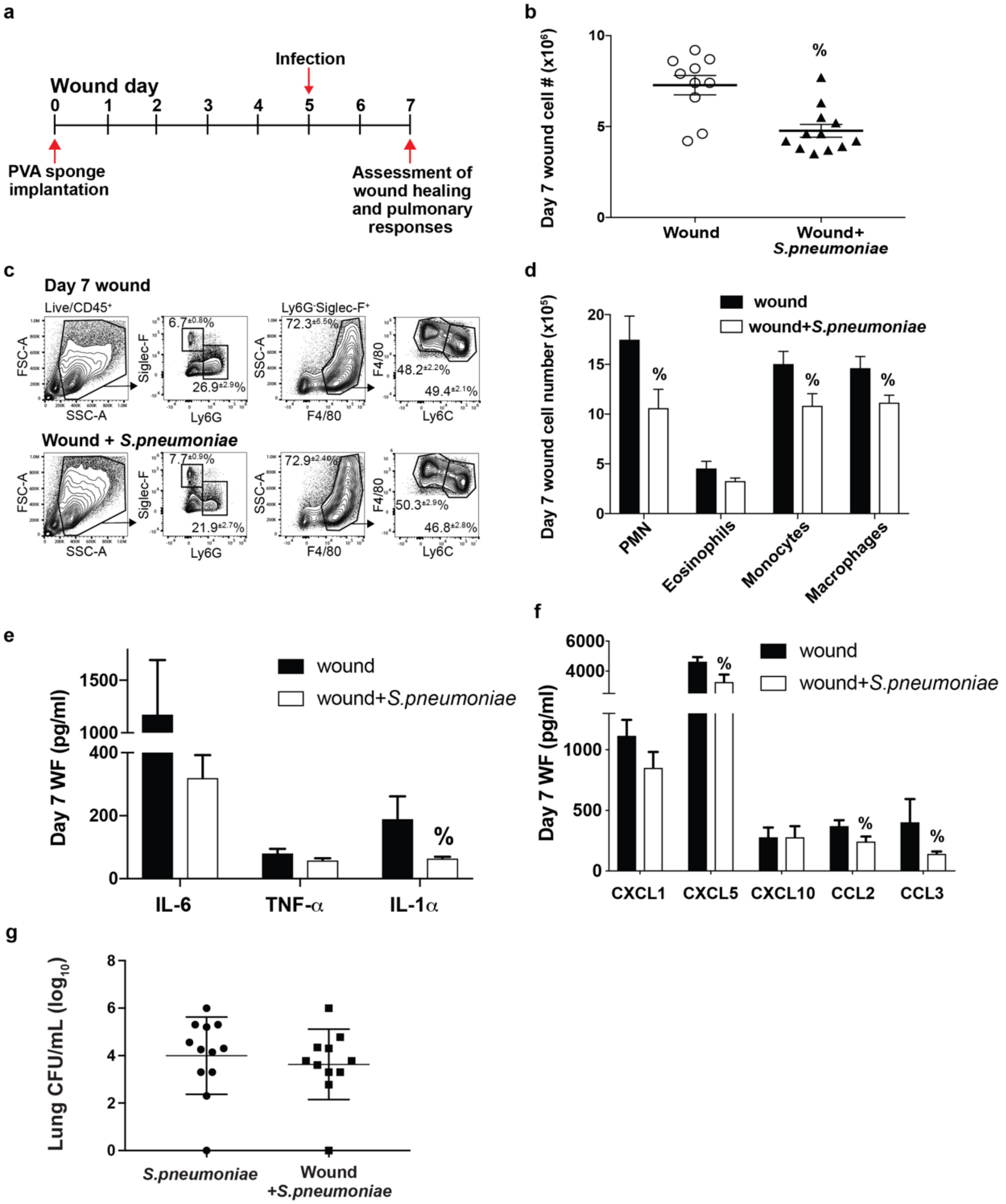


**Figure S6. The effect of pulmonary *Streptococcus pneumoniae* infection on cutaneous wound healing.** Mice were wounded by the subcutaneous implantation of PVA sponges, then infected intranasally with $5 \times 10^6$ CFU *S. pneumoniae* 5 days later. Wound cellular and cytokine responses were assessed on wound day 7 (a). The onset of pulmonary *S. pneumoniae* infection suppressed the overall cellularity of the wound on day 7 (b). The frequency of innate leukocyte subsets infiltrating the wound on day 7 was not altered by *S. pneumoniae* infection (c), although the absolute number of neutrophils (PMN), monocytes, and macrophages was suppressed by the pulmonary infection (d). The effect of pulmonary infection on wound fluid (WF) concentrations of the proinflammatory cytokines IL-6, TNF-α, and IL-1α (e) and the chemokines CXCL1, CXCL5, CXCL10, CCL2, and CCL3 (f) was determined. Prior wounding did not alter the lung bacterial burden as assessed on wound day 7 (48h post-infection) (g). Data are the mean ± SEM with a minimum n=10. % indicates p≤0.05 comparing wound+*K. oxytoca* to wound groups.

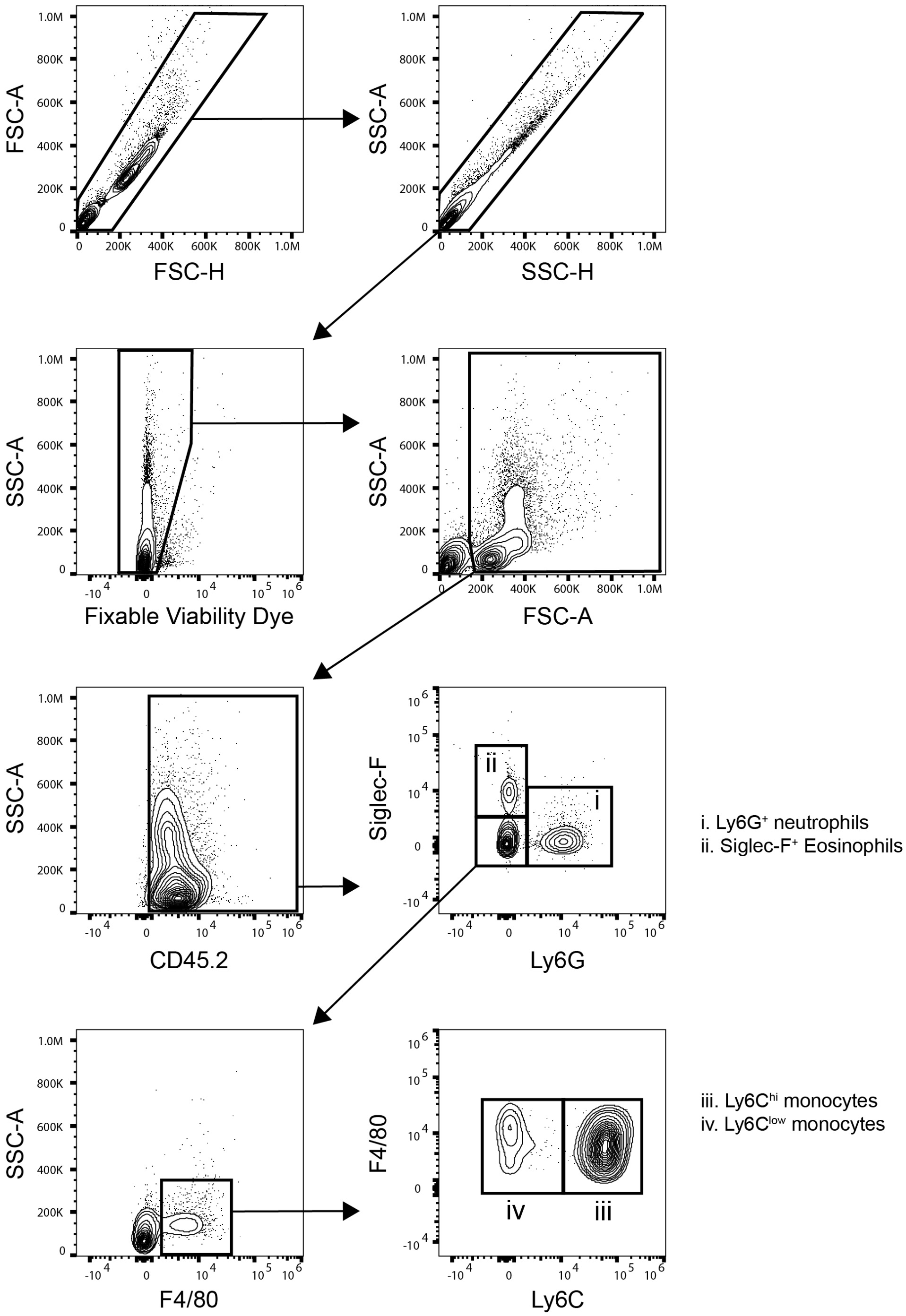


**Figure S7. Representative gating strategy to identify blood innate leukocyte populations.** This gating strategy was employed to quantify innate leukocytes in the blood. Doublets were excluded, then dead cells were removed from the analysis using a fixable viability dye. Cell debris and residual red blood cells were excluded by size using the FSC-A and SSC-A parameters. Hematopoietic cells were identified as CD45.2$^{+}$. Neutrophils (i) were identified as Ly6G$^{+}$Siglec-F$^{-}$. Eosinophils (ii) were identified as Siglec-F$^{+}$Ly6G$^{-}$. F4/80$^{+}$ monocytes were gated from the Ly6G$^{-}$Siglec-F$^{-}$ population. F4/80$^{+}$ cells were fractionated into Ly6C$^{hi}$ inflammatory monocyte (iii) and Ly6C$^{low}$ patrolling monocyte populations (iv).

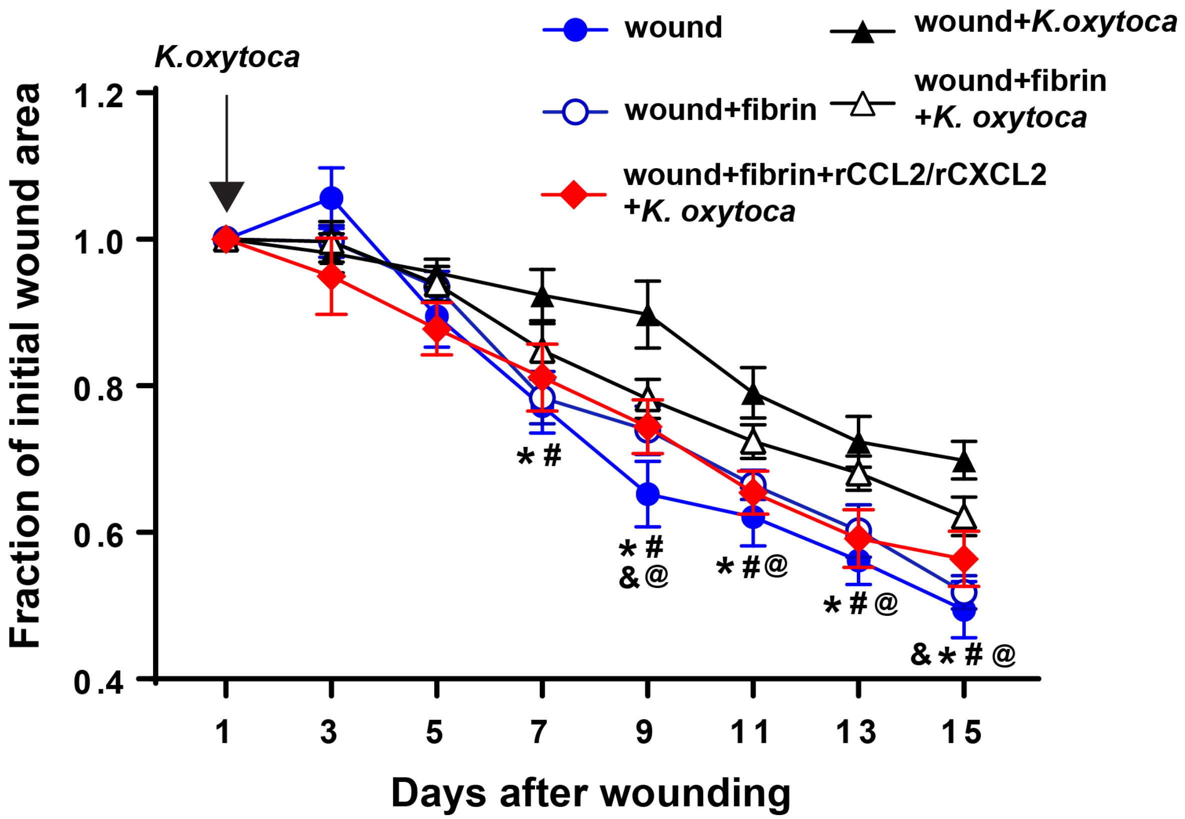


**Figure S8. The effect of fibrin sealant supplemented with recombinant chemokine on excisional tail wound healing in mice with pulmonary *K. oxytoca* infection.** Mice were wounded by tail skin excision. A cohort of wounded mice was infected intranasally with *K. oxytoca* one day later. Wounds remained untreated, were treated with fibrin sealant (Tisseel), or were treated with fibrin sealant supplemented with recombinant CCL2 and CXCL2 as described in Materials & Methods. The effect of all treatments on the rate of tail wound closure is presented. Fibrin treatment alone did not significantly alter the rate of tail wound closure in infected mice, whereas the combination of fibrin sealant and recombinant CCL2+CXCL1 significantly accelerated wound closure beginning on wound day 9. * indicates a statistically significant change between wound (uninfected) and wound+*K. oxytoca*, # indicates a statistically significant change between wound+fibrin and wound+*K. oxytoca*, & indicates a statistically significant change between wound and wound+fibrin+*K. oxytoca*, @ indicates a statistically significant change between wound+*K. oxytoca* and wound+fibrin+rCCL2/rCXCL1+*K. oxytoca* groups.
